# R-Loop control and mitochondria genome stability require the 5’-3’ exonuclease/flap endonuclease OEX1

**DOI:** 10.1101/2024.11.05.621957

**Authors:** Déborah Schatz-Daas, Anaïs Le Blevenec, Fabio G. Moratti, Kin Pan Chung, Pierre Mercier, Rana Khalid Iqbal, Elody Vallet, André Dietrich, Ralph Bock, Frédérique Weber-Lotfi, José M. Gualberto

## Abstract

Maintenance of the plant organelle genomes involves factors mostly inherited from their bacterial symbiotic ancestors. In bacteria, a major player in genome maintenance is DNA Polymerase I (Pol I), which provides a 5’-3’-exonuclease/flap-endonuclease activity required for multiple replication and repair functions. In plant organelles, DNA polymerases POL1A and POL1B are evolutionarily derived from DNA Pol I but lack this domain. In Arabidopsis, OEX1 and OEX2 (Organellar Exonucleases 1 and 2) represent this missing domain and are targeted to mitochondria and chloroplasts, respectively. An *oex1* mutant allele shows developmental and fertility defects that correlate with the differential segregation of mitochondrial DNA (mtDNA) subgenomes generated by recombination, suggesting that OEX1 processes replication and recombination intermediates whose accumulation results in genome instability. Alternative splicing generates two OEX1 isoforms that can differentially interact with POL1A and POL1B and variably affect mtDNA repair.

Recombinant OEX1 has 5’-3’-exonuclease and flap endonuclease activities, the latter being a key function in replication and repair. Furthermore, OEX1 has high affinity for RNA:DNA hybrids, rapidly degrading RNA in Okazaki-like structures and R-loops. Consistent with a role in suppressing R-loops, *oex1* plants accumulate R-loops in highly transcribed mtDNA regions. Taken together, our results show that OEX1 plays multiple important roles in the processes required to maintain mtDNA stability.

## Introduction

Mitochondria and chloroplasts have their own genomes (mtDNA and cpDNA), which are remnants of their past autonomous prokaryotic lifestyle. The mtDNA contains a limited number of genes (37 in humans and 50-60 in plants), yet these genes are crucial for cell survival, as they code for essential subunits of oxidative phosphorylation complexes or are required for their expression (Knoop, 2004; Chevigny et al., 2020). Although plant and animal mtDNA code for a similar number of genes, plant mitogenomes are considerably larger, with sizes ranging from 200 kb up to 11 Mb (Sloan et al., 2012), and have been observed in diverse structural and organizational forms, including linear, circular, and branched configurations (Oldenburg and Bendich, 1996; Backert et al., 1997; Manchekar et al., 2006). This high variability in size and structure results from rearrangements through homologous recombination (HR) involving repeated sequences, which are abundant in plant mitogenomes (Kubo and Newton, 2008). These repeated sequences, ranging in size from several kilobases to less than 100 bp, engage in HR with different frequencies according to their size and genomic position, thus leading to the highly dynamic structure of land plant mitogenomes (Small et al., 1989; Hanson and Bentolila, 2004). Recombination is also the main driving force for the rapid evolution of the plant mtDNA organization, which can significantly differ between closely related species and even among different accessions of the same species (Mower et al., 2007; Fertet et al., 2021).

Despite its remarkable organizational plasticity, plant mtDNA protein-coding sequences remain remarkably conserved across most species, indicating the presence of highly effective repair mechanisms. Both base excision repair (BER) and HR-dependent repair processes occur in plant mitochondria (Boesch et al., 2009; Ferrando et al., 2018; Trasvina-Arenas et al., 2018). In particular, break-induced replication (BIR) and illegitimate recombination through microhomology-mediated BIR (MMBIR) have also been proposed as important processes driving replication, repair, and evolution of plant mitogenomes (Cappadocia et al., 2010; Christensen, 2013; Gandini et al., 2023). Genetic mutations that impair HR efficiency or specificity can significantly affect plant fitness, resilience to stress, and fertility (Gualberto and Newton, 2017).

Factors already identified as involved in plant mitochondrial HR include RecA-like recombinases, several different types of single-strand DNA (ssDNA)-binding proteins, a MutS-like protein (MSH1) involved in the rejection of homoeologous recombination, and the branch-migration helicases RECG and RADA (Chevigny et al., 2020). Importantly, all HR-dependent repair processes require a DNA polymerase, and while the animal mtDNA is replicated by the single enzyme DNA Pol γ, the maintenance of land plants mitochondrial and plastid genomes involves two multifunctional DNA polymerases derived from prokaryotic DNA Pol I (POL1A and POL1B). Both are dually targeted to plastids and mitochondria (Mori et al., 2005; Carrie et al., 2009) and differentially participate in replication and repair processes (Parent et al., 2011; Peralta-Castro et al., 2020). However, many other factors that should play pivotal roles in plant mtDNA replication and repair, including key nucleases, have yet to be identified. During replication, such nucleases should be involved in the processing of Okazaki-like primers prior to the ligation of the newly synthesized DNA segments. Primer removal could occur by strand displacement by a DNA polymerase, via the generation of a flap structure to be processed by a flap endonuclease (FEN) (Zheng and Shen, 2011; Ma et al., 2022), or by an RNase H (Randall et al., 2019). Both mechanisms can coexist, as in animal mitochondria, where the flap endonuclease FEN1, RNase H1, endonuclease DNA2, and the ssDNA-specific nuclease MGME1 play roles in potentially redundant pathways for primer removal (Uhler and Falkenberg, 2015). Exonucleases and RNase H can additionally be involved in the control of R-loops, which are three-stranded structures generated during transcription, in which an RNA segment is hybridized to one of the DNA strands (Roy and Lieber, 2009; Ginno et al., 2012). If not regulated, R-loops and unprocessed Okazaki primers can lead to replication fork stalling, genome rearrangements, and increased recombination (Gan et al., 2011; Aguilera and Garcia-Muse, 2012; Wimberly et al., 2013). Additionally, a FEN or 5’-3’ exonuclease should be involved in plant mitochondrial repair pathways, for strand resection following double-strand breaks and the processing of recombination intermediates. A FEN may also play a role in long-patch BER, to process the flap formed by strand displacement during DNA synthesis (Ranalli et al., 2002). To date, no flap endonuclease or functional analogs of MGME1 or DNA2 have been identified in plant organelles. A source of exonuclease and FEN activities could be the 5’-3’ nuclease domain of a bacterial-type DNA polymerase Pol I, a domain essential for bacterial Pol I repair functions. However, plant organellar POL1A and POL1B lack a 5’-3’ exonuclease domain.

Here we describe OEX1, an Arabidopsis mitochondrial nuclease conserved in the entire plant lineage and phylogenetically related to the 5’-3’ exonuclease domain of prokaryotic DNA Pol I. OEX1 is essential for mtDNA stability, and its loss leads to severe plant growth defects. *In vitro*, OEX1 degrades dsDNA processively from 5’ to 3’ and shows efficient flap endonuclease activity, capable of processing substrates generated by the displacement activity of DNA polymerase. We show that OEX1 can degrade RNA in Okazaki-like RNA:DNA hybrids, and that *oex1-1* mutants accumulate R-loops within highly transcribed mtDNA regions. OEX1 exists as two isoforms resulting from alternative splicing, which have similar substrate specificities *in vitro*, but differ in processivity, affinity to the two organellar DNA polymerases, and differentially affect mtDNA segregation and repair. Our results show that OEX1 is an exonuclease/flap endonuclease that potentially plays multiple roles in mtDNA maintenance.

## Results

### Two Arabidopsis genes code for potential organellar exonucleases

The plant organellar DNA polymerases POL1A and POL1B lack the N-terminal 5’-3’ exonuclease domain that in bacterial Pol I plays important roles in DNA repair and the maturation of Okazaki fragments during replication (Lovett, 2011). In bacteria, the loss of this activity results in conditional lethality, whereas mutations just affecting the DNA polymerase or the 3’-5’ exonuclease domains of DNA Pol I are viable (Lehman and Uyemura, 1976). A BLAST search of the Arabidopsis proteome for potential counterparts to the 5’-3’ exonuclease domain of DNA Pol I revealed two candidate genes, At3g52050 and At1g34380, which we preliminarily named *OEX1* and *OEX2*, respectively, for <u>O</u>rganellar <u>Ex</u>onuclease 1 and 2 (Figure 1A). They correspond to the genes previously identified through similar searches conducted by other researchers (Sato et al., 2003; Moriyama and Sato, 2014). The encoded proteins contain a distinctive PIN-like 5’ nuclease domain and an H3TH (helix-3-turn-helix) DNA-binding domain, a structure also found in the 5’-3’ exonuclease domain of type A DNA polymerases, in eukaryotic flap endonuclease FEN1 and in RNase H of bacteriophage T4 (Matelska et al., 2017). Phylogenetic analysis revealed that OEX1 is present in all groups of the green lineage, including land plants and green algae, as previously observed (Moriyama and Sato, 2014). In contrast, OEX2 is found exclusively in land plants (Figure 1B). The similarity between OEX1 and OEX2 is lower than that between OEX1 and the orthologous bacterial sequences (Figure 1C). Additionally, none of the intron positions is shared between the *OEX1* and *OEX2* genes. This suggests that *OEX2* is either a paralog of *OEX1* that has undergone rapid divergence, or was acquired independently of *OEX1* later in evolution, for example, through horizontal gene transfer.

**Figure 1.**
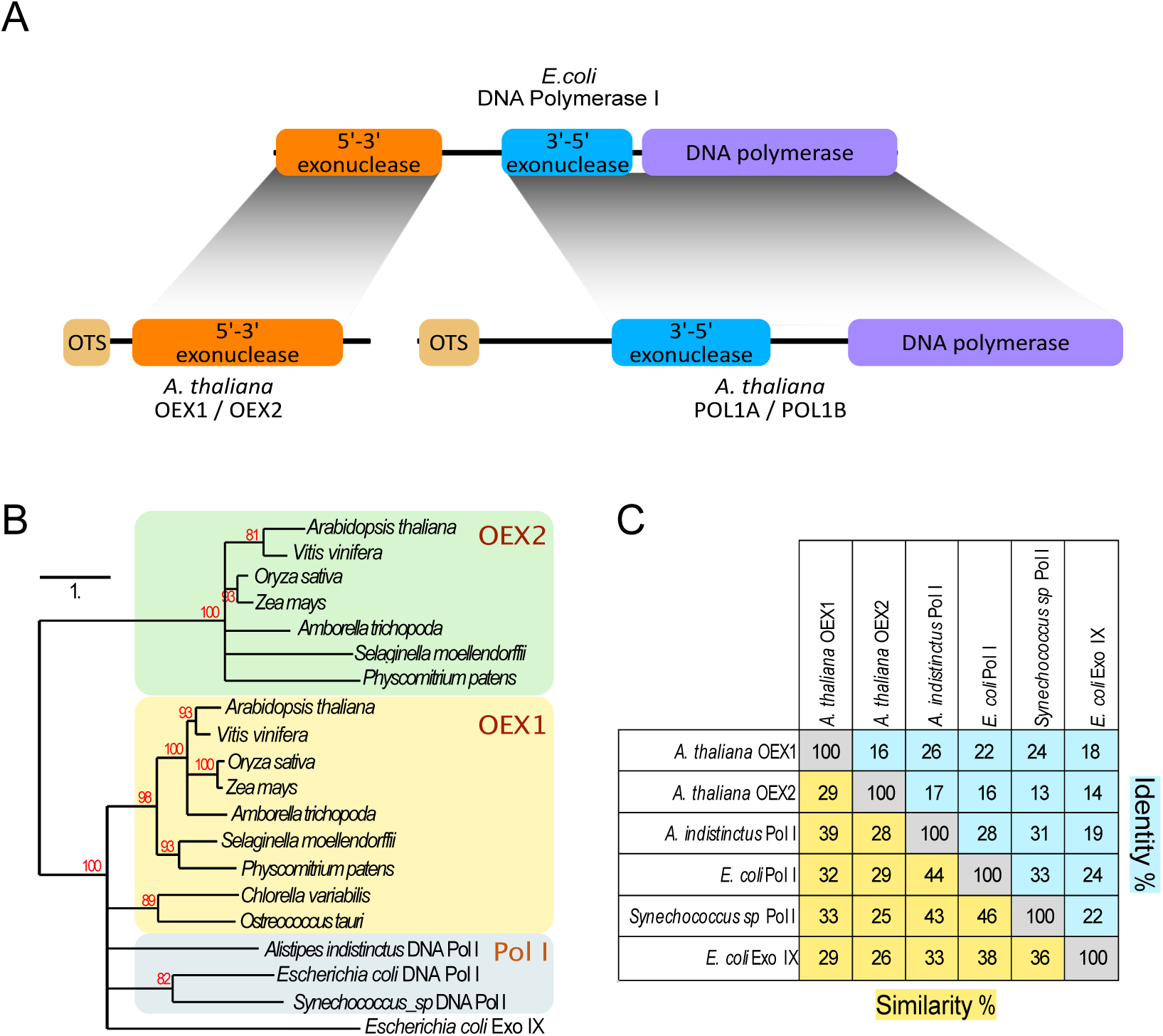
OEX1 and OEX2 are plant organellar orthologs of bacterial exonucleases. **A)** Two Arabidopsis genes encode putative organellar proteins similar to the 5’-3’ exonuclease domain of bacterial DNA polymerase I, a domain absent from plant organellar DNA polymerases POL1A and POL1B (OTS: organellar targeting sequence). **B)** Phenogram showing that OEX1 orthologs are present across the green lineage, including land plants and green algae, while OEX2 orthologs are restricted to land plants. **C)** Similarity between OEX1 and OEX2 compared to the similarities to bacterial proteins. The data suggest that the two plant genes did not diverge from a common ancestor and instead, may have distinct origins.

### OEX1 is targeted to mitochondria and OEX2 to chloroplasts

Both OEX1 and OEX2 display N-terminal extensions compared to their bacterial homologs. Predictions suggest that these extensions are mitochondrial and chloroplast targeting sequences, respectively (https://services.healthtech.dtu.dk/services/TargetP-2.0/; https://urgi.versailles.inra.fr/predotar/). To confirm the *in silico* predictions, genetic constructs encoding the fusion proteins OEX1-GFP and OEX2-GFP were introduced in wild-type Arabidopsis via Agrobacterium-mediated transformation. Stably transformed Arabidopsis plants expressing OEX1-GFP revealed a typical mitochondrial localization (Figure 2A). This was confirmed with an OEX1-mCherry construct introduced into a mitochondrial marker line (mt-GFP), by the co-localization of the mCherry and GFP signals. Interestingly, the mCherry fluorescence was not evenly distributed within the mitochondrial matrix but accumulated in distinct foci. DAPI staining showed that these foci co-localize with DNA, suggesting that OEX1 is recruited to mitochondrial nucleoids (Figure 2B). Attempts to express OEX2-GFP in stably transformed Arabidopsis plants were unsuccessful, presumably because of transgene silencing, but transient expression in leaf epidermal cells showed strict chloroplast localization (Figure 2C). In maize, the ortholog of OEX2 was found significantly enriched in the nucleoid proteome of plastids (Majeran et al., 2012).

**Figure 2.**
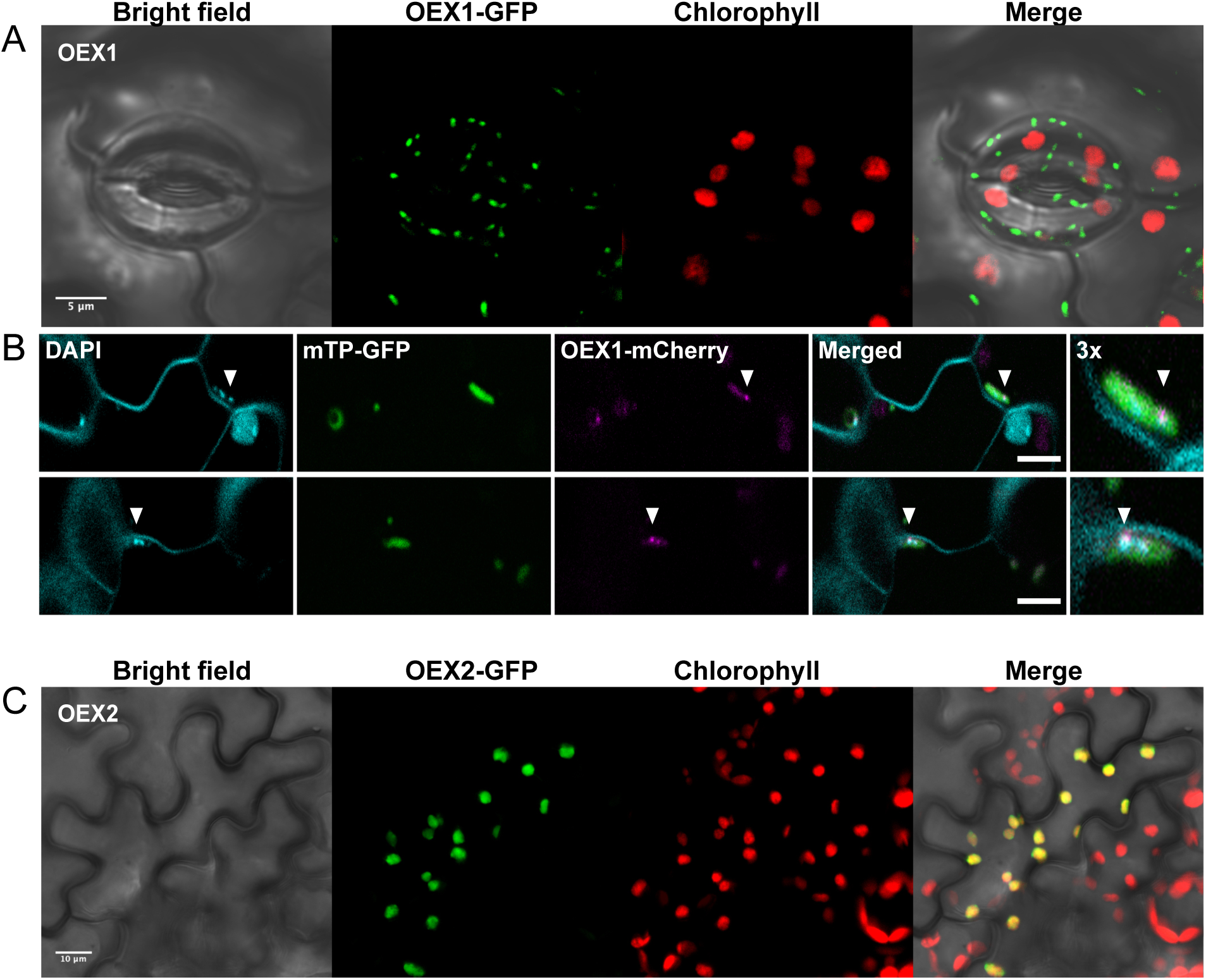
OEX1 and OEX2 are targeted to mitochondria and chloroplasts, respectively. **A)** Mitochondrial targeting of OEX1:GFP stably expressed in transgenic Arabidopsis plants. **B)** OEX1 is localized in mitochondrial nucleoids, as revealed by colocalization of OEX1-mCherry with DAPI-stained nucleoids (mT-GFP: mitochondrially targeted GFP serving as compartment-specific marker protein). **C)** Chloroplast targeting of OEX2:GFP in *Nicotiana benthamiana* leaf epidermal cells following biolistic transfection.

### OEX1 is required for normal plant development and fertility

In our subsequent work, we focused on characterizing the mitochondrial OEX1. The developmental and spatial expression patterns of OEX1 were studied by expressing the ß-glucuronidase (GUS) gene under the control of the *OEX1* promoter. GUS staining was observed in cotyledons and very young developing leaves, as well as in the central vascular cylinder of roots, in young pistils and stamens (Figure S1). No expression was detected in gametophytic tissues such as pollen and ovules. These results are mostly consistent with published RNAseq data, which show that OEX1 is predominantly expressed in young leaves, petals, and pistils (Klepikova et al., 2016).

The consequences of *OEX*1 loss were analyzed in a T-DNA insertion mutant. Searching publicly available collections, several candidate mutants were initially identified, of which we ultimately could validate a single line (GABI_911E05). This mutant line, containing an insertion within the gene, was designated *oex1-1*. The T-DNA insertion in *oex1-1* was mapped 10 nucleotides downstream of the ninth exon of *OEX1*. RT-PCR showed that the last six exons are no longer transcribed, a result that was confirmed by RNAseq analysis (Figure S2). The mutation disrupts the protein-coding sequence at the level of the PIN-like nuclease active site, and results in the loss of 152 out of 425 amino acids, including the entire H3TH DNA-binding domain, indicating that *oex1-1* is a true knock-out line. Homozygous *oex1-1* plants are viable, but severely impaired in development (Figure 3A). Compared to wild-type (WT) plants, they display stunted growth, distorted leaves and flowers, and have very low fertility. Although *oex1-1* plants can produce inflorescences, flowers are abnormal, with curved pistils, and pollen production is visibly reduced. The small flower stalks carry small siliques that mainly contain aborted embryos (Figure 3A), and of the few mature seeds obtained many failed to germinate. The observed phenotype is fully penetrant, affecting all homozygous *oex1-1* plants, although with some variation in the severity among individual T2 plants (Figure 3A). In subsequent generations, the phenotypes worsened, and third-generation homozygous mutants were very strongly affected in growth and unable to flower. Reciprocal backcross attempts with WT plants were unsuccessful, suggesting that both male and female *oex1-1* gametes are affected. Similar phenotypes have been described for mutants of several genes involved in mtDNA maintenance. In particular, the *oex1-1* phenotypes are reminiscent of those of *radA* plants, deficient in the RADA branch migration helicase (Chevigny et al., 2022).

**Figure 3.**
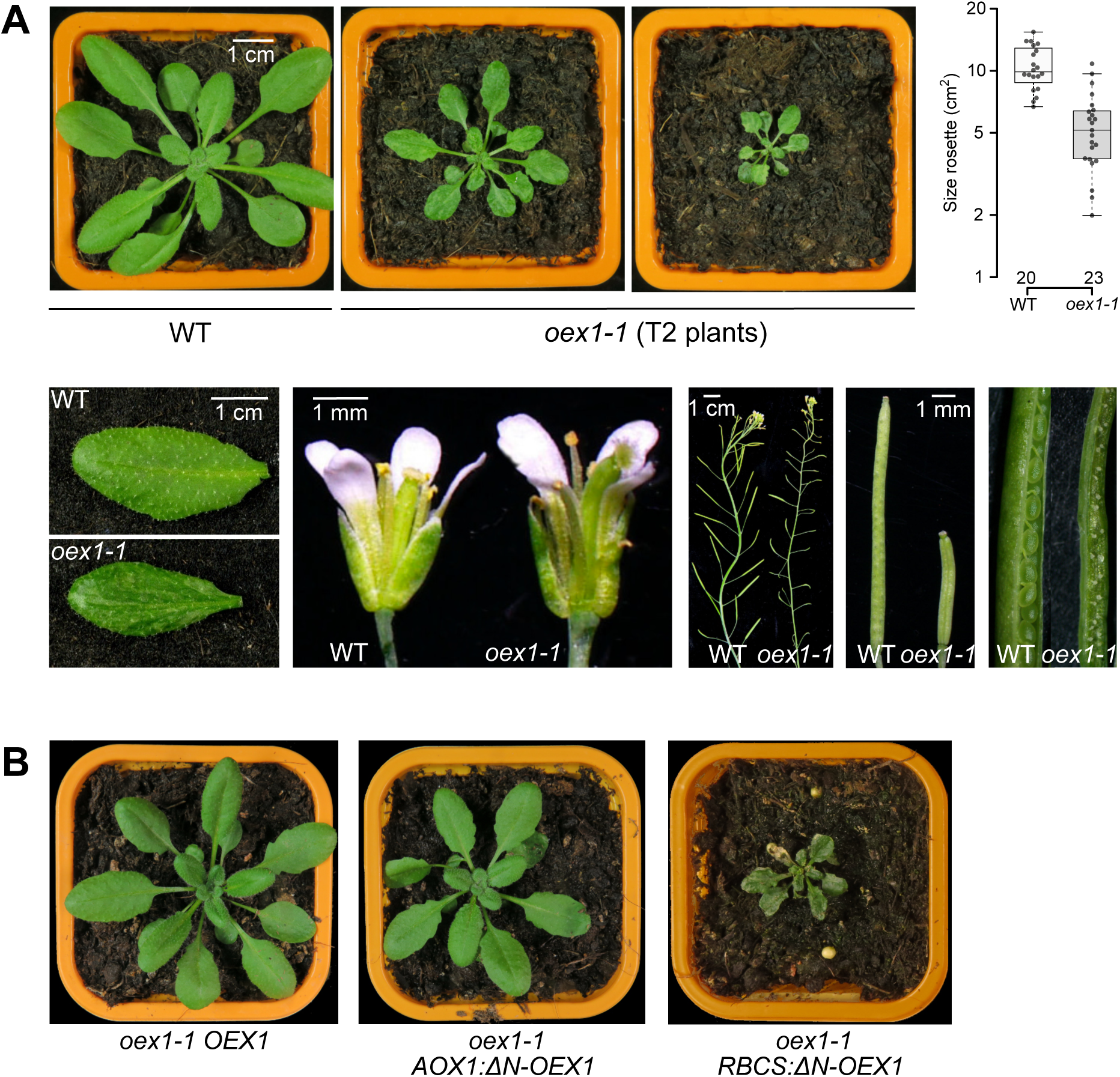
Loss of OEX1 has severe effects on plant growth and development. **A)** Representative images of 4-week-old wild-type (WT) and plants, showing growth retardation, distorted leaves with chlorotic sectors, flower deformities, and reduced fertility of the mutant. The severity of the phenotypes can vary significantly among individual plants within the first homozygous mutant generation (T2 plants), as highlighted in the box plots showing the high variation in rosette size in *oex1-1*. Flowers harbor only few pollen grains that adhere to the papillae of the stigma. Siliques are small and mainly contain aborted embryos. **B)** The developmental phenotypes of the *oex1-1* mutant are fully complemented in *oex1-1* plants expressing OEX1 under its native promoter (*oex1-1 OEX1* plant) and in mutants expressing OEX1 targeted exclusively to mitochondria, as achieved by fusion to the targeting sequence of AOX1 (*oex1-1 AOX1:ΔN-OEX1* plant). However, chloroplast-specific OEX1 expression by fusion to the targeting sequence of RBCS does not rescue the phenotype (*oex1 RBCS:ΔN-OEX1* plant).

To explore possible changes in mitochondrial morphology, *oex1-1* rosette leaf sections were examined by transmission electron microscopy (TEM). The TEM images showed that mitochondria in *oex1-1* mesophyll cells were smaller and more electron-dense than in WT plants of similar size. In contrast, the morphology of chloroplasts was unaffected (Figure S3).

The deleterious mutant phenotypes were fully complemented by expression of the *OEX1* cDNA under the native *OEX1* promoter. Complemented plants (*oex1 OEX1*) were phenotypically normal and indistinguishable from WT plants (Figure 3B). To confirm that the deleterious phenotype was caused by lack of OEX1 activity in mitochondria only, and not by the possible presence of undetectably low protein levels in chloroplasts, hemicomplementation experiments were also conducted. To this end, *oex1-1* plants were transformed with constructs designed to specifically target OEX1 to either mitochondria or chloroplasts (see Methods). As shown in Figure 3B, full complementation of the mutant phenotype was achieved when OEX1 was specifically targeted to mitochondria, whereas no complementation was observed when OEX1 was solely targeted to chloroplasts. Together, these results support the hypothesis that the mutant phenotype of *oex1-1* plants is primarily due to defects in mitochondria.

### OEX1 plays a critical role in maintaining mitochondrial genome stability

Mutants deficient in factors involved in mtDNA maintenance and repair exhibit genome instability, mostly due to the accumulation of cross-over products resulting from recombination mediated by small repeated sequences (100-500 bp) (Gualberto and Newton, 2017). To test whether lack of OEX1 results in cross-over product accumulation, we performed total DNA sequencing (Illumina) on *oex1-1* vs. Col-0 plants. For this analysis, *oex1-1* seedlings from the first homozygous mutant generation, segregating from a heterozygous line, were used. The relative abundance (stoichiometry) of the different genomic regions of the mtDNA and cpDNA were evaluated, according to the coverage profiles, to highlight enriched or depleted regions.

The profile of the plastid genome sequence coverage from *oex1-1* was indistinguishable from that of the WT, thus providing no evidence of cpDNA instability (Figure 4B). In stark contrast, the mtDNA profile of *oex1-1* plants was significantly perturbed, showing positive and negative stoichiometric changes, in sharply demarcated, discrete regions of the genome (Figure 4A). Several of those discrete genomic regions in the mtDNA are flanked by directly oriented repeats (e.g., repeats A (556 bp), L (249 bp), and EE (127 bp), according to the nomenclature of repeats annotated in the mtDNA genome of Col-0, accession BK010421). This suggests that the mtDNA stretches corresponding to these regions may represent subgenomic molecules generated by a “loop-out” mechanism during recombination, a phenomenon observed in other mtDNA recombination mutants (Wallet et al., 2015; Chevigny et al., 2022). To confirm this hypothesis, the accumulation of cross-over products resulting from recombination involving repeats L, F, and EE was tested by qPCR, as described before (Miller-Messmer et al., 2012) and according to the cartoon at the top of Figure 4C. These repeats were selected because they are consistently mobilized in other mtDNA repair mutants and are small enough for qPCR analysis. As expected, a significant increase in cross-over products was observed compared to WT levels (Figure 4C). An asymmetric accumulation of primarily one type of cross-over was seen, consistent with previous findings in other mutants with mtDNA recombination defects (Gualberto and Newton, 2017). This asymmetry is likely due to the involvement of the error-prone break-induced replication (BIR) repair pathway (Christensen, 2013; Gualberto and Newton, 2017; Kockler et al., 2021).

**Figure 4.**
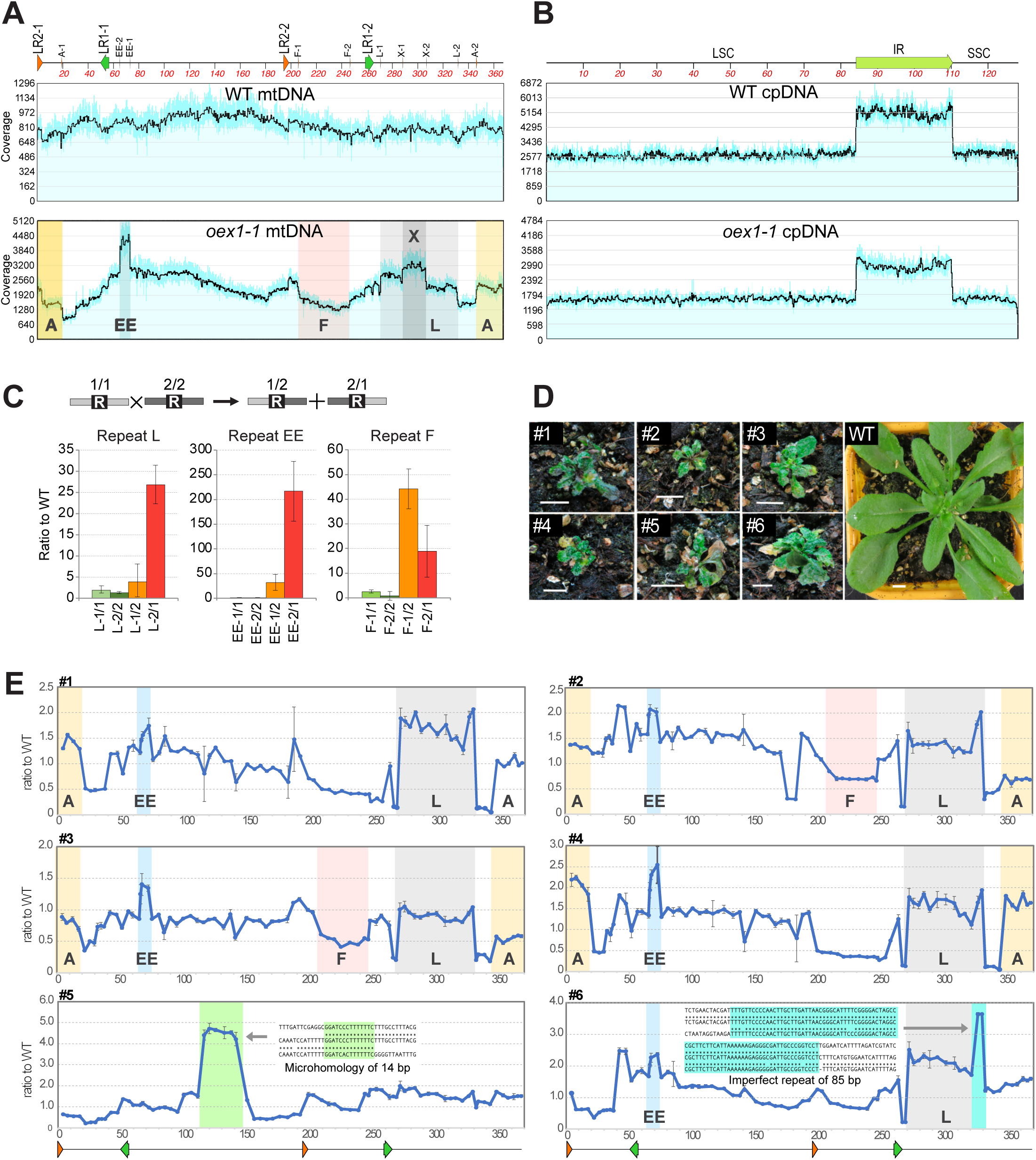
Structural alterations of the mtDNA in *oex1* mutants. **A)** Sequence coverage of the mtDNA in *oex1-1* vs. WT, revealing regions with altered stoichiometry flanked by direct repeats. The coordinates are those of the Col-0 mtDNA sequence. The positions of the mtDNA repeats LR1 and LR2 are shown on top of the plotted area. Regions with changed stoichiometry are shadowed and annotated according to the repeat name (repeats A, F, L, X and EE, of 556 bp, 350 bp, 249 bp, 204 bp and 127 bp, respectively). **B)** Plastid genome analysis as performed in A), showing no effects of OEX1 loss on cpDNA stability. LSC: large single copy region, SSC: small single copy region, IR: inverted repeat (only one copy is included). **C)** qPCR confirmation of the accumulation in *oex1-1* plants of recombination products across repeats L, F and EE. The parental sequences 1/1 and 2/2, and the corresponding crossover products 1/2 and 2/1 were quantified by qPCR, as schematically represented on top of the figure for the sequences flanking a repeated sequence “R”. Error bars correspond to *SD* values from three biological replicates. **D** and **E)** Analysis of the relative copy numbers of the different mtDNA regions in individual severely affected T4 generation *oex1* plants. Sequences spaced about 5 kb apart in the mtDNA were quantified by qPCR. Regions with altered stoichiometry flanked by direct repeats are shadowed as in A. In plants #5 and #6, regions that are highly increased in copy number correspond to subgenomes resulting from recombination involving a microhomology of 14 bp and an imperfect repeat of 85 bp, respectively. The parental and recombined sequences are shown.

To assess the extent of mtDNA changes in individual *oex1-1* plants, the relative copy number of different mtDNA regions was measured by qPCR in six *oex1-1* T3-generation plants that displayed severe growth and developmental impairments (Figure 4D). All plants showed marked changes in the stoichiometry of different mtDNA regions, though the extent of these changes varied greatly among plants (Figure 4E). Most changes occurred in the regions flanked by repeats A, L, F, and EE (plants #1 to #4 in Figure 4E), as it had been seen in T2-generation plants. However, some plants (plants #5 and #6) showed a particularly large increase in copy number of sequences not associated with any annotated repeat. Further investigation revealed that these regions corresponded to subgenomic molecules formed through illegitimate recombination (i.e., involving sequences of little or imperfect homology), with one involving an 85 bp imperfect repeat and the other a 14 bp microhomology (Figure 4E). These results suggest that, in the absence of OEX1, illegitimate recombination activities are activated, which can recruit small repeats and microhomologies. Consequently, the mitochondrial genome stoichiometry is altered, as the newly formed subgenomes seem to behave as episomes and are preferentially amplified (through unknown mechanisms). A similar scenario was described previously for the *recG1* and *radA* mutants (Wallet et al., 2015; Chevigny et al., 2022). Thus, the severe growth phenotype of *oex1-1* plants correlates with defects in mtDNA maintenance, resulting in the accumulation of ectopic recombination products and significant alterations in the stoichiometry of the different mtDNA regions. However, the mtDNA regions with lower copy numbers in *oex1-1* plants do not contain any identified gene(s) that could explain the profound developmental defects observed in these plants.

### Two OEX1 isoforms are generated by alternative splicing

RT-PCR amplification and sequencing of *OEX1* transcripts from Arabidopsis leaves revealed two mRNA variants caused by alternative splicing of the 6^th^ exon. This splicing was found in 6 out of 11 cDNA clones sequenced, and is supported by publicly available EST sequences (https://www.arabidopsis.org). The two mRNA variants encode proteins that differ by 18 amino acids, which we named OEX1a and OEX1b (Figure 5A).

**Figure 5.**
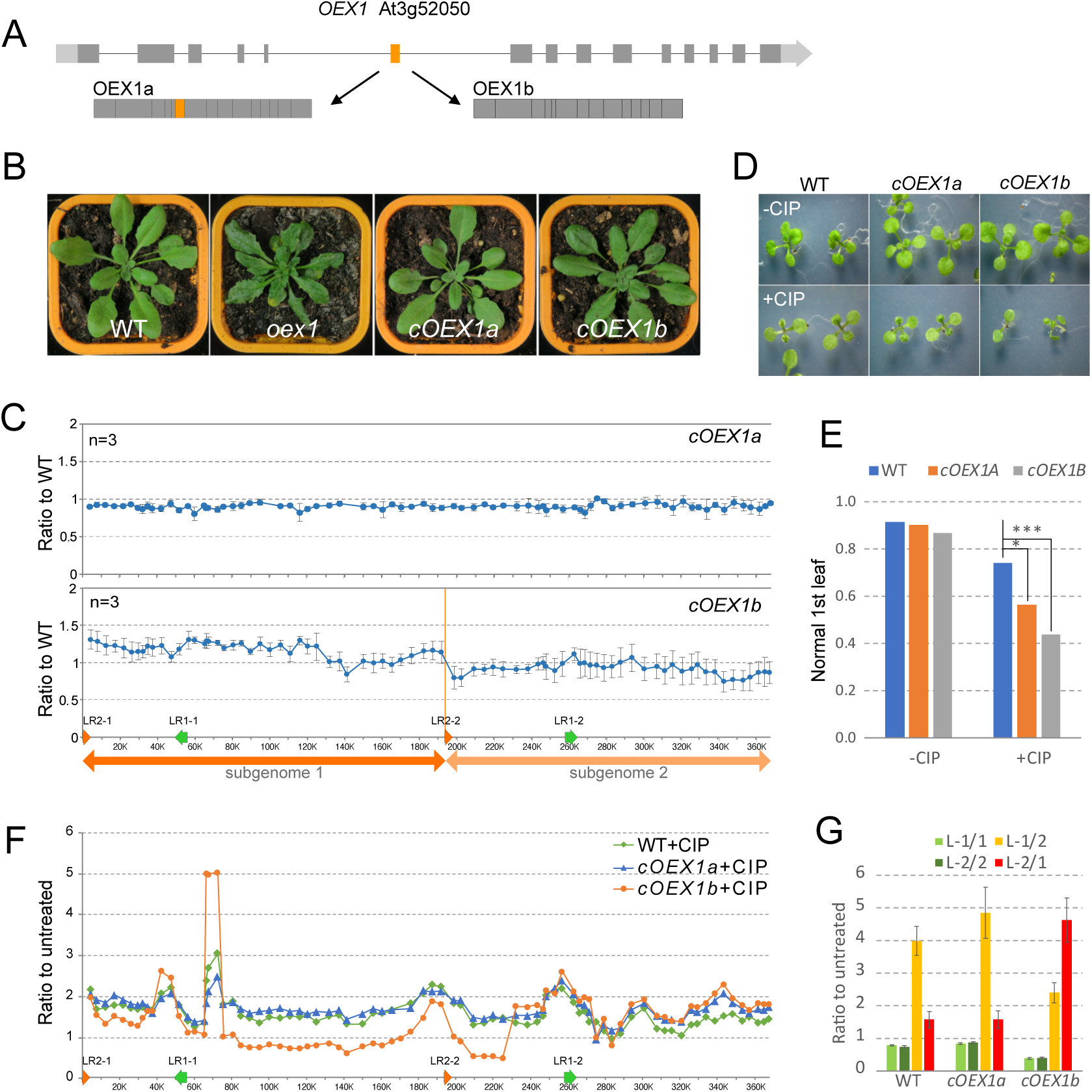
Alternative splicing results in the expression of two OEX1 isoforms. **A)** Schematic representation of the two proteins resulting from alternative splicing of exon 6. **B)** Complementation of the *oex1-1* mutant with the OEX1a or OEX1b coding sequences (*cOEX1a* and *cOEX1b* plants, respectively) under the control of the endogenous *OEX1* promoter. Both constructs complemented the severe developmental phenotypes of *oex1-1*. **C)** Analysis of the mtDNA of *cOEX1a* and *cOEX1b* plants by qPCR, as described in Figure 4E, showing full complementation by *cOEX1a*, but retention of a slightly remodeled mtDNA in *oex1 cOEX1b*, with altered copy number of the subgenomes defined by large repeat LR2. Results represent the average and *SD* of three biological replicates. **D** and **E)** Phenotypes of plants grown under genotoxic stress induced by 0.5 µM ciprofloxacin (CIP). Plants complemented with *cOEX1b* were more sensitive, as assessed by the percentage of plants still able to develop normal first true leaves, at a CIP concentration that had only a mild effect on the WT. Asterisks indicate statistically significant differences according to the **χ^2^** test (*: p<0.05, **: p<0.01). **F)** Effect of CIP treatment on the stability of the mtDNA in WT and *cOEX1a* and *cOEX1b* plants. Analysis by qPCR, as in C. **G)** Relative accumulation of crossover products resulting from recombination involving pair of repeats L, as described in Figure 4C, in CIP-treated plants as compared to the non-treated control. The crossover product L-1/2 accumulates in CIP-treated WT and complemented plants *cOEX1a*, while it is the reciprocal product L-2/1 that accumulates in *cOEX1b*.

The 6^th^ exon-encoded extension retained in the OEX1a isoform is conserved across all flowering plants at the DNA level, though its sequence conservation is low (Figure S4A). Corresponding transcript variants have been identified, among others, in soybean (XM_006574959.4), grapevine (<u>XM_010652395.3</u>), maize (<u>XM_008660501.4</u>), rice (<u>XM_015776708.3</u>) and Amborella (<u>XM_020676298.1</u>). However, this extension is absent from OEX2 and from the 5’-3’ exonuclease domain of bacterial DNA Pol I (Figure S4A). Structural modeling suggests that the extension is located near the active site and the H3TH DNA-binding domain (Figure S4B), potentially influencing OEX1 activity and/or substrate specificity. The relative abundance of the *OEX1*a and *OEX1b* transcripts in different plant tissues was assessed by RT-qPCR, using variant-specific primers (Figure S4C). Both mRNA forms were detected at similar levels in most tissues analyzed, except for flower buds, where the ratio of *OEX1b* to *OEX1a* was higher, suggesting that OEX1 may be developmentally regulated through alternative splicing (Figure S4C).

The impact of the two OEX1 isoforms on mtDNA maintenance was further examined by complementing the *oex1-1* mutant with constructs harboring either OEX1a or OEX1b cDNAs, under the control of the OEX1 promoter. Heterozygous *oex1-1* plants were transformed with the constructs, self-pollinated, and homozygous *oex1-1* plants containing the transgene were analyzed. Both isoforms successfully complemented the development defects of *oex1-1*, and the resulting plants (*cOEX1a* and *cOEX1b*) were indistinguishable from WT (Figure 5B). However, at the molecular level, while stoichiometric mtDNA replication was fully restored to wild-type levels in *cOEX1a* plants, *cOEX1b* plants showed a slight imbalance in the relative abundance of two mtDNA subgenomes flanked by large repeat 2 (LR2, 4.2 kb in size; Figure 5C). The complemented *cOEX1a* and *cOEX1b* plants were further challenged for repair capacity by growth under genotoxic stress induced by ciprofloxacin (CIP), as described previously (Schatz-Daas et al., 2022). At the diagnostic concentration of 0.5 µM CIP, most WT plants were still able to develop normal first true leaves, whereas *cOEX1b* plants and, to a lesser extent, *cOEX1a* plants either failed to develop their first true leaves or these leaves were mostly wrinkled and distorted (Figure 5D and 5E). qPCR analysis of mtDNA copy number across the genome revealed that CIP treatment significantly affects WT plants, causing significant changes in the relative stoichiometry of different mtDNA regions. A similar effect was observed in CIP-treated *cOEX1a*-complemented plants. However, more pronounced changes were seen in *cOEX1b* plants (Figure 5F). We further analyzed the accumulation of the crossover products from recombination involving repeat L, which previous studies showed to accumulate differently in CIP-treated WT and repair-deficient mutants (Schatz-Daas et al., 2022). As described, in CIP-treated WT plants, sequence L-1/2 accumulates, likely due to HR-mediated repair of DNA breaks. A similar effect was observed in *cOEX1a* plants, but it was much reduced in *cOEX1b* plants, where the crossover sequence L-2/1 accumulated instead (Figure 5G). Thus, defects in the replication, repair, and segregation of mtDNA subgenomes were not fully restored in either of the complemented plants, particularly not in *cOEX1b*. This suggests that OEX1a and OEX1b are two isoforms that are only partially redundant with respect to their roles in mtDNA maintenance.

### OEX1a and OEX1b may differentially interact with the organellar DNA polymerases

Because of the sequence similarity between OEX1 and the 5’-3’ exonuclease domain of type A bacterial DNA polymerases, a yeast two-hybrid (Y2H) assay was used to test whether OEX1 can interact with the plant organellar DNA polymerases POL1A or POL1B to reconstitute a presumed holoenzyme. No interaction was observed when the OEX1a or OEX1b coding sequences were fused to the GAL4 activating domain (AD) and POL1A or POL1B to the binding domain (BD). However, in the reverse configuration, both BD-OEX1a and BD-OEX1b interacted with AD-POL1A, while only BD-OEX1b interacted with AD-POL1B (Figure S5A and S5B). These results further suggest different *in vivo* roles for OEX1a and OEX1b. Because POL1A and POL1B were described as having differential roles in repair (Parent et al., 2011), the different effects of OEX1a and OEX1b on mitochondrial DNA stability might be causally related to their differential association with POL1B.

Yet, proteomic data revealed that OEX1 migrates as a single band in blue-native gels, much smaller than 100 kDa (https://complexomemap.de/; (Senkler et al., 2017); Figure S5C). The apparent size matches that of monomeric OEX1 (about 43 kDa) rather than that of a complex with POL1A or POL1B (both of which are larger than 100 kDa). Furthermore, quantitative proteomic analysis indicated that OEX1 is more abundant than the DNA polymerases, with about 40 molecules of OEX1 per mitochondrion compared to 5 and 2 molecules of POL1A and POL1B, respectively (Fuchs et al., 2019). These data rather support the idea that OEX1 does not form permanently stable complexes with the DNA polymerases. Instead, the interaction (suggested by the Y2H assays) may be transient.

### Loss of OEX1 has no apparent detrimental effect on mitochondrial transcripts

To assess whether mtDNA instability in *oex1-1* affects mitochondrial gene expression, mitochondrial transcript abundances in *oex1-1* and WT Col-0 were analyzed by RT-qPCR, as described previously (Baudry et al., 2022). For genes with introns, the integrity of the exon-exon junctions in the mRNAs was also examined. *oex1-1* exhibited an increase in the abundance of most transcripts, with only a few transcripts showing a modest decrease. No splicing defects were detected (Figure 6). Because the reshuffling of mtDNA sequences in *oex1-1* could potentially impact transcript processing, the RNA-seq coverage profiles from WT Col-0 and *oex1-1* were comparatively inspected. This analysis revealed no apparent changes in the processing of mitochondrial protein-coding transcripts (Figure S6). This finding suggests that the severe growth defects in *oex1-1* plants are not caused by deficient expression of a specific mitochondrial gene. Interestingly, several gene transcripts, including *atp1*, *rps4*, *rps7*, *ccmC*, *matR*, *ccmFN1* and *ccmFN2,* were more than two-fold higher in *oex1-1*, according to RT-qPCR. In certain cases, this increase may be attributable to the increased gene copy number. For example, *ccmC*, *ccmFN1* and *ccmFN2,* which are clustered in the subgenome defined by repeat L, or *atp1* that is encoded by the EE episome that is amplified at the DNA level in the mutant. In plants complemented with either *OEX1a* or *OEX1b*, mitochondrial transcript abundances were restored to WT levels (Figure 6).

**Figure 6.**
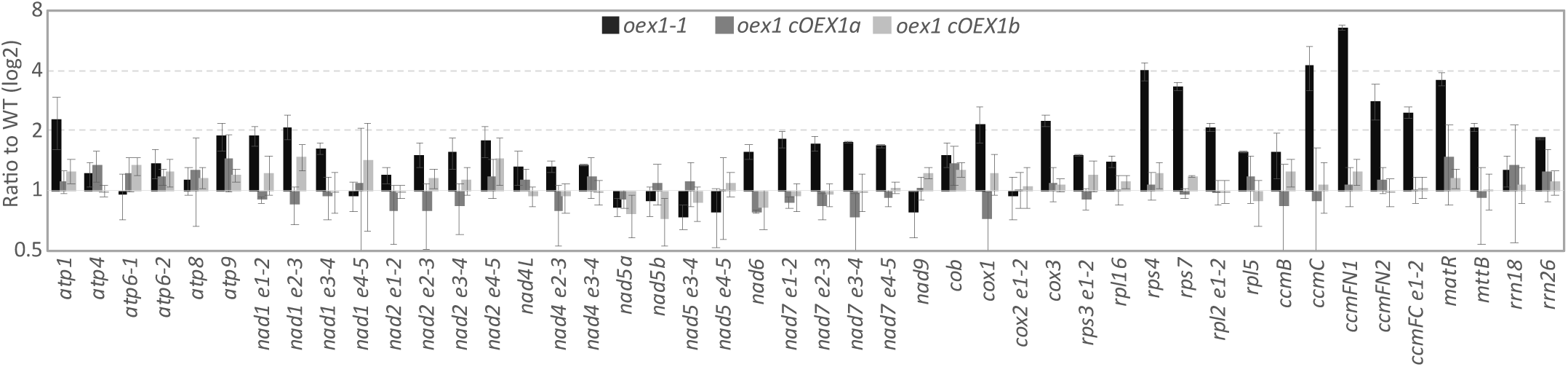
Accumulation of mitochondrial transcripts in *oex1* mutant plants. **A)** RT-qPCR analysis of steady-state levels of mitochondrial protein-coding transcripts and rRNAs in *oex1-1* plants and the complemented lines *cOEX1a* and *cOEX1b.* Results were normalized to a set of nuclear housekeeping genes. The exon-exon borders were quantified for genes containing introns, to assay for possible splicing defects. Results are displayed at log2 scale and represent the average and *SD* from three biological replicates.

### OEX1 is a 5’-3’ exonuclease/flap endonuclease

To investigate the enzymatic activities of the OEX1 nuclease, recombinant OEX1a and OEX1b proteins were expressed in *E. coli* and purified (Figure S7). According to the known structure of PIN-like nucleases, it was predicted that residue D250 is part of the active site. Therefore, a putative non-catalytic mutant protein version (D250A) was also designed and purified (Figure S7B).

Preliminary experiments indicated that recombinant OEX1a was capable of degrading 5’-labeled linear dsDNA, which was therefore used as a substrate to optimize reaction conditions for maximal OEX1a activity (Figure S8). OEX1a efficiently degraded a 25-mer dsDNA substrate, showing strict 5’-3’ exonuclease activity, without evidence of endonuclease activity, as no intermediate products were detected with 5’-labeled substrates (Figure 7A). To confirm that this activity is attributable to OEX1a and does not result from bacterial proteins contaminating the purified protein, the D250A non-catalytic mutant version was purified and tested under identical conditions, and showed no activity. Under the same conditions OEX1b also showed 5’-3’ exonuclease activity, but with less processivity than OEX1a, and a significant portion of the substrate remained intact after 30 minutes incubation (Figure 7B). The degradation pattern of a 3’-labeled dsDNA substrate ([GTCA]_5_GTCCC) suggested sequence-dependent distributive activity, with pausing at G:C base pairs (Figure 7A and B). This was confirmed by testing substrates with 1, 2 or 3 consecutive Gs, which showed intermediate degradation products whose ends mapped to the G positions, with the strongest signal for the G_3_ substrate (Figure S9A).

**Figure 7.**
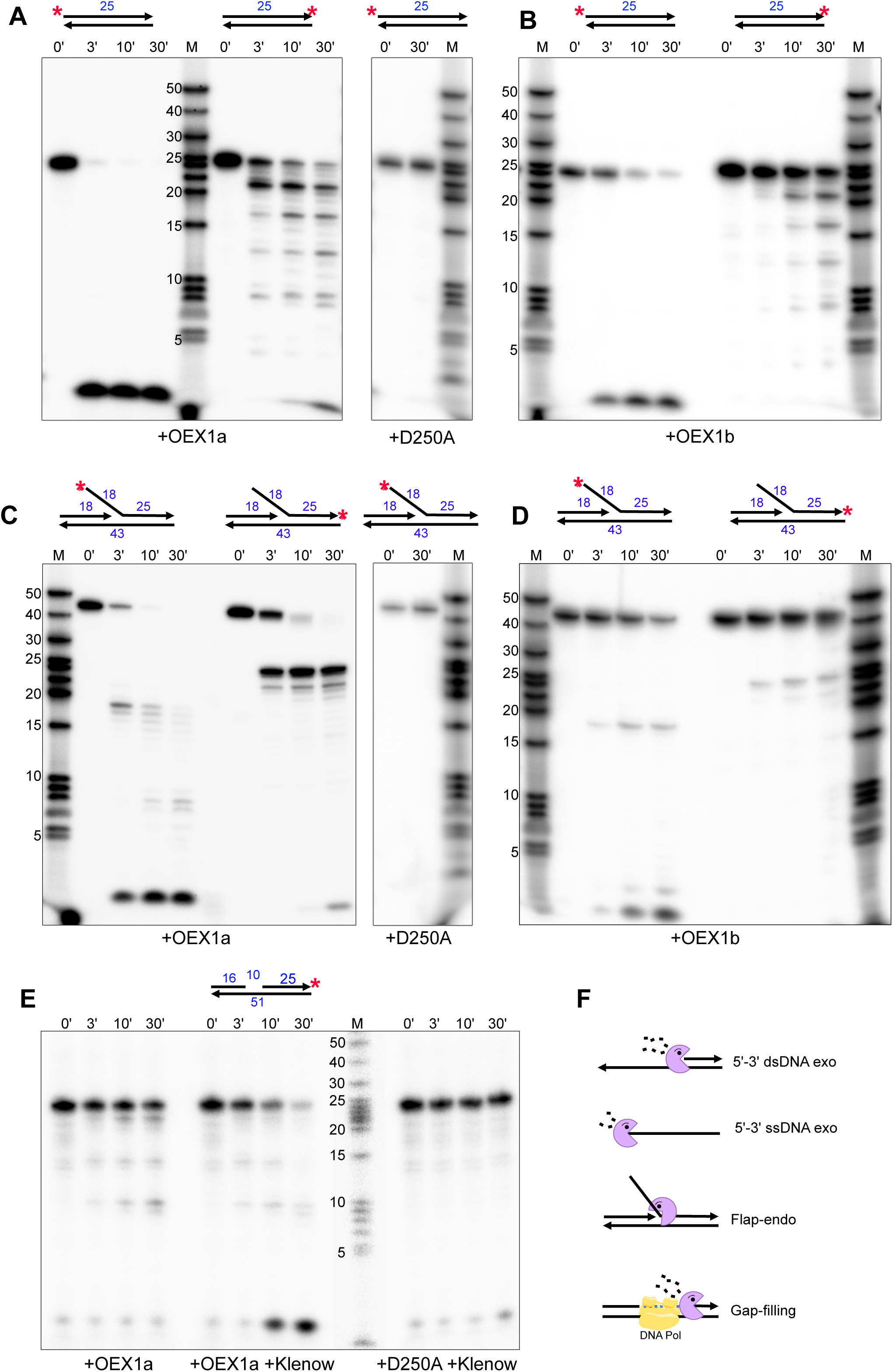
Exonuclease and flap endonuclease activities of OEX1. **A)** Analysis of the exonuclease activity of the wild-type OEX1a protein and the non-catalytic mutant D250A on a dsDNA substrate, schematically represented above the gel images and labeled either at the 5’ or the 3’ end (red stars). Substrates and degradation products were analyzed by electrophoretic separation in denaturing polyacrylamide gels. Size of DNA markers are indicated in nucleotides **B)** Analysis as in A, but for isoform OEX1b. **C)** Analysis as in A, but with a substrate mimicking flap structures. **D)** Analysis as in C, but for OEX1b. **E)** Analysis as in A, but for a substrate with a 10 nt gap, in the absence or presence of a DNA polymerase (Klenow enzyme). **F)** Summary of OEX1 activities on the various DNA substrates tested.

To test for flap endonuclease activity, a 5’ flap-like substrate was prepared and labeled either at the 5’ or 3’ end of the flap strand. The flap strand was specifically designed without sequence homology to the complementary DNA, to prevent the formation of alternative structures. Precise processing of the flap was expected to produce either an 18 or a 25 nucleotide-labeled product, depending on the labeling site. Both OEX1a and OEX1b, but not the non-catalytic mutant, could precisely process the flap (Figure 7C,D). Similar to the 5’-3’ exonuclease activity, OEX1a showed higher processivity on the flap substrate than OEX1b.

Following flap cleavage by OEX1a, the resulting 5’-labeled 18-mer was further degraded, implying that OEX1 can also degrade ssDNA. This prompted us to test OEX1 activity on ssDNA. Both OEX1a and OEX1b showed 5’-3’ exonuclease activities, as evidenced by using a 3’-labeled substrate (Figure S9B). However, these activities were much weaker than those observed on dsDNA, and most of the ssDNA substrate remained intact after 30 minutes. As for dsDNA, OEX1b showed much weaker activity than OEX1a. Testing a 5’-labeled ssDNA substrate further revealed that OEX1a also has weak 3’-5’ activity, removing 1–2 nucleotides before the label was removed by the 5′-3′ activity (Figure S9B). This low activity on ssDNA *in vitro* may not be biologically relevant, or alternatively, could play a role in the surveillance of unscheduled replication triggered by small ssDNA fragments.

Next, we tested whether OEX1 could play a role in base excision repair (BER), namely whether it can recognize single-strand gaps on dsDNA (mimicking BER intermediates) and extend them by removing a few nucleotides. On dsDNA substrates with gaps of 1, 3 or 10 nucleotides, OEX1a showed minimal exonuclease activity. Although it could extend the gaps regardless of their size (Figure S9C), this occurred with much lower processivity than the activity at the 5′ end of dsDNA, suggesting that OEX1 alone is inefficient in extending single-strand gaps. To test if gap extension is improved by strand displacement by a DNA polymerase, the experiment was repeated with the Klenow enzyme added after OEX1a. Under these conditions, gaps were processed significantly faster, indicating that OEX1a activity on gapped dsDNA is enhanced by strand displacement (Figure 7E). No significant activity was observed with Klenow polymerase and the D250A non-catalytic mutant, confirming that the results cannot be explained by a contaminant or the 3’-5’ exonuclease activity of the Klenow enzyme. In summary, our *in vitro* assays show that OEX1 has 5’-3’ exonuclease activity on dsDNA, flap endonuclease activity, and gap-extension capability in cooperation with the strand displacement activity of a DNA polymerase (Figure 7F).

### OEX1a readily degrades the RNA moiety in RNA:DNA hybrids and is involved in the processing of R-loops

Given the potential role of OEX1 in the processing of Okazaki fragments, its activity on RNA:DNA hybrids was tested. Of a 3’-labeled RNA paired to a complementary DNA primer (RNA:DNA substrate), OEX1a rapidly degraded the RNA strand (Figure 8A), with apparently much higher processivity than on dsDNA substrates (as seen in the 3’-labeled product in Figure 7A). On a substrate mimicking an Okazaki-like RNA primer on the lagging DNA strand, consisting of a dsDNA region followed by a small gap and an RNA:DNA duplex, OEX1a also rapidly degraded the RNA oligonucleotide (Figure 8B). This contrasts with the weak activity of OEX1a on a similar substrate made of DNA duplexes only, where strand displacement by a DNA polymerase was needed to accelerate the reaction (Figure 7E). Finally, when OEX1a was tested on single-stranded

**Figure 8.**
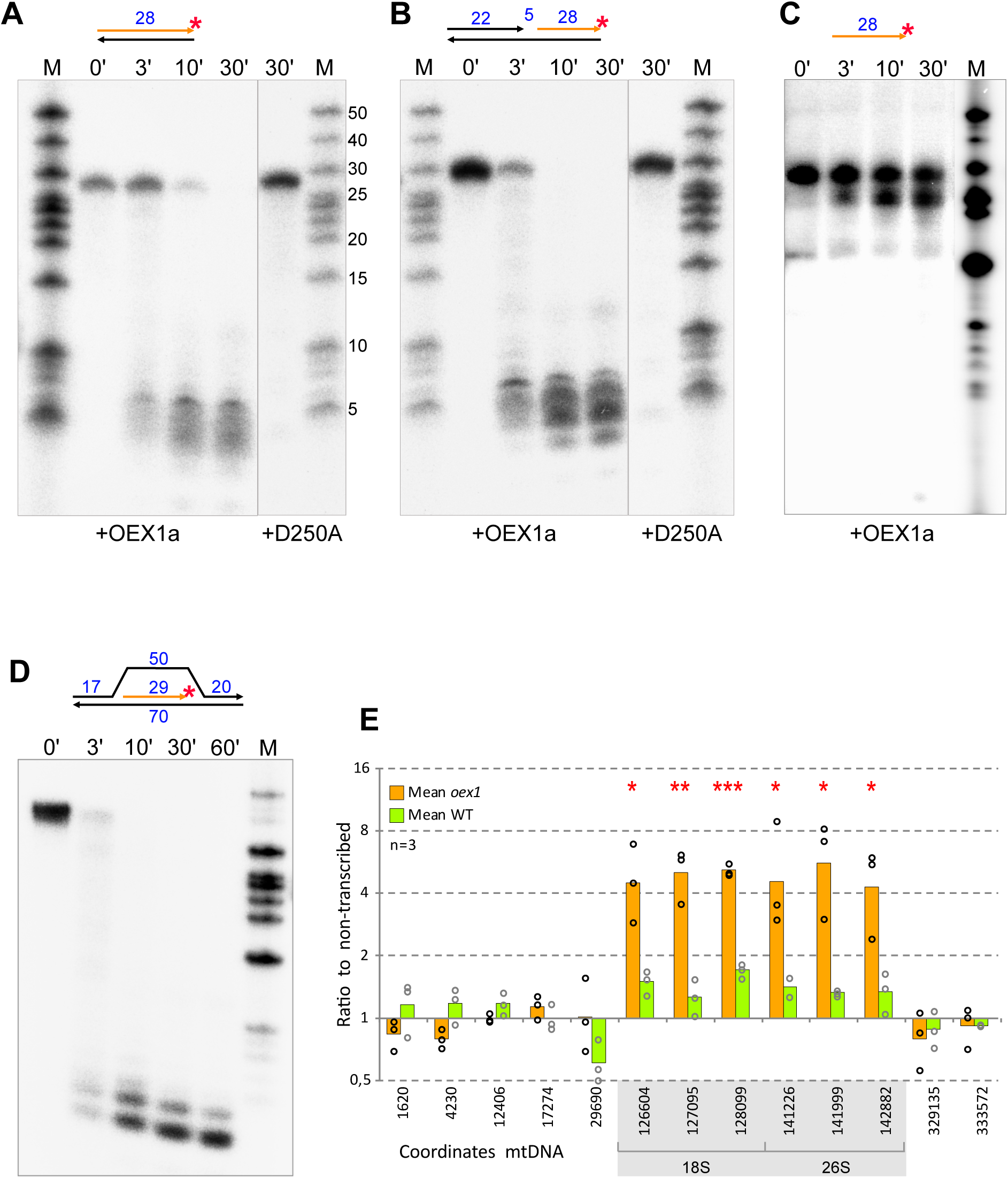
OEX1 exonuclease activity on RNA:DNA hybrids, and *in vivo* contribution to the clearance of R-loops in the mtDNA. **A)** Analysis of the exonuclease activity of wild-type OEX1a protein and the non-catalytic mutant D250A on a RNA:DNA hybrid labeled at the 3’ end of the RNA strand (red star). Substrates and degradation products were analyzed by electrophoresis in denaturing polyacrylamide gels. **B)** Assay as in A, but with a substrate representing an Okazaki-type RNA fragment. **C)** Assay as in A, showing that OEX1 has no activity on a single-stranded RNA substrate. **D)** Assay as in A, showing that OEX1 has high activity on the RNA moiety of an R-loop. **E)** DRIP-qPCR analysis of R-loops in *oex1* plants compared to the WT. Results are the average of three biological replicates. The data show an increase of R-loops in the *oex1* mutant, in the mtDNA region corresponding to the highly transcribed 18S and 26S rRNAs. Results were normalized against non-transcribed regions of the mtDNA. Asterisks represent statistically significant changes by Student’s t-test (*: p<0.05, **: p<0.01, ***: p<0.001).

RNA, no significant nuclease activity was detected (Figure 8C). In summary, the *in vitro* assays show that RNA:DNA hybrids are apparently the preferred substrates of OEX1. *In vivo,* such substrates may be formed during replication (e.g., Okazaki fragments) or might be by-products of transcriptional activities (e.g., R-loops). Both are known to compromise genome stability if unresolved (Crossley et al., 2019). We, therefore, explored the potential involvement of OEX1 in resolving R-loops. First, we tested the *in vitro* activity of OEX1a on a substrate mimicking an R-loop, with the 5’ dsDNA ends biotinylated to protect them from OEX1 activity. Upon incubation with OEX1a, the RNA component of this R-loop substrate was promptly degraded (Figure 8D). Next, we assessed whether *oex1-1* plants accumulate R-loops, by DNA:RNA immunoprecipitation (DRIP). To this end, nucleic acids from WT and *oex1-1* plants were immunoprecipitated using the S9.6 antibody specific for RNA:DNA hybrids, and the precipitated DNA was analyzed by qPCR. As also reported by others (Sanz and Chedin, 2019), the signal-to-background ratio in the DRIP experiments was low. Nevertheless, a clear and significant increase was observed in the *oex1-1* samples compared to the WT (Figure 8E). This overaccumulation of R-loops could be linked to the mtDNA instability observed in *oex1-1* mutant plants.

## Discussion

### Origin of the OEX1 and OEX2 genes and their relationship to bacterial counterparts

Due to its symbiotic origin, mtDNA replication and stability involve a combination of factors, many of which are derived from the prokaryotic ancestors of organelles. In eubacteria, a 5’-3’ exonuclease activity is essential for cell viability. It is provided by either Pol I or ExoIX orthologs (e.g., YpcP in *B. subtilis*) (Fukushima et al., 2007). These enzymes have a structure-specific endonuclease activity that cleave at the junction between a 5′ single strand and a duplex. The DNA polymerase I activity at a nick displaces the DNA upstream of the enzyme, forming a 5′ flap that is subsequently cleaved by the 5′ endo/exonuclease (Anstey-Gilbert et al., 2013; Randall et al., 2019). In many bacterial species, the two enzymes co-exist, but their respective contributions to DNA replication and genome stability remain unclear. A recent study found that they have different substrates affinities, hinting at specialized roles during replication (Lowder and Simmons, 2023). Still, either enzyme is sufficient for cell survival, whereas double mutation leads to synthetic lethality (Fukushima et al., 2007). Thus, in bacteria lacking ExoIX activity, the 5’-3’ exonuclease activity of Pol I is indispensable (Diaz et al., 1992). Conversely, some bacterial genera have only an ExoIX ortholog and lack a Pol I-like repair polymerase. They correspond to class 2 in Fukushima’s classification, and they are all symbiotic or infectious bacteria, including the probable symbiotic ancestor of mitochondria. It is thus tempting to speculate that plant mitochondria fall into the class 2 systems and that OEX1/2 originated from a bacterial ExoIX. However, phylogenetic analysis suggests that OEX1/2 are closer to the N-terminal domain of Pol I than to ExoIX, implying that plant OEX1/2 evolved from a Pol I that lost its polymerase domain. The phylogenetic analysis also indicates that OEX2 is not a paralog of OEX1, as their similarity to each other is lower than that to bacterial orthologs, suggesting distinct acquisition events.

Our subcellular localization analyses strongly suggest that OEX1 is exclusively mitochondrial and specifically localized in nucleoids, consistent with a role in mtDNA metabolism. In agreement with the confocal microscopy results, hemicomplementation of *oex1-1* plants with a construct targeting OEX1 exclusively to mitochondria restored the WT phenotype, confirming that OEX1 is active only in mitochondria. Similarly, the exclusive chloroplast localization of OEX2-GFP is in line with the identification of the maize OEX2 ortholog in the plastid nucleoid proteome (Majeran et al., 2012). Taken together, these results suggest that OEX1 and OEX2 fulfill equivalent nuclease functions in mitochondria and plastids.

### Major potential role of OEX1 in mtDNA replication

To date, details of how mtDNA replicates have been elucidated in only a few model systems. In animals, replication of the small and compact mtDNA is highly specialized. It requires mitochondrial RNase H1 to generate RNA primers by cleaving longer transcripts within R-loops stabilized by ssDNA-binding protein mtSSB (Falkenberg et al., 2024). After replication, the RNA primers are removed by RNase H1 and nuclease MGME1, which cuts the ssDNA flaps formed by the displacement activity of mitochondrial DNA polymerase γ (Uhler et al., 2016). Other nucleases that are dually targeted to both the nucleus and mitochondria such as FEN1 and DNA2 may also be involved.

In plants, however, there is no MGME1 ortholog, and mtDNA replication and maintenance rely on many prokaryotic-like factors. The processing of RNA primers bound to the mtDNA during replication likely mirroqqqqrs the bacterial system, with OEX1 emerging as the primary candidate enzyme for this function. The marked FEN and gap-extension activities of recombinant OEX1, facilitated by DNA polymerase, support the hypotheses that OEX1 has a central role in cleaving the small flaps formed by strand displacement when DNA polymerase reaches a downstream 5’-end. Consistent with this predominant role in replication, the recombinant protein efficiently recognized and processed the RNA strand of an RNA:DNA hybrid *in vitro*, as well as structures mimicking Okazaki fragments in a replication fork (Figure 8). In these experiments, OEX1 was more processive on RNA than on DNA substrates, and was processive on gap-containing double-stranded DNA only in the presence of DNA polymerase activity. These results suggest that the elimination of RNA sequences hybridized to DNA is the main role of OEX1. Therefore, maturation of plant mitochondrial Okazaki fragments by OEX1 could occur either via exonucleolytic degradation and/or via FEN activity on the flaps previously formed by strand displacement by DNA polymerase. Interestingly, bacterial YpcP also showed its highest activity on RNA:DNA duplexes, recognizing Okazaki fragments and flap substrates (Randall et al., 2019). Organellar RNase H1 enzymes may contribute to the removal of replication primers. However, in plants, the loss of mitochondrial RNase H1B alone does not show adverse effects on plant development, a finding that is in stark contrast to the vital importance of mitochondrial RNase H1 in animals (Cheng et al., 2021; Misic et al., 2022).

### Other potential roles of OEX1 in the mtDNA maintenance

HR is the primary mechanism for repair and maintenance of plant mitogenomes, and requires several nucleases. A 5’-3’ exonuclease is typically required for strand resection at double-strand breaks. In bacteria, this function is handled by the RecBCD complex or by RecJ in the RecFOR pathway, both of which are highly processive (Kowalczykowski, 2015). OEX1 has 5’-3’ exonuclease activity on blunt-ended dsDNA, but the *in vitro* tests showed that its activity on these substrates is distributive rather than processive, suggesting that it is unlikely to be directly involved in strand resection. Nonetheless, it seems possible that OEX1 functions *in vivo* in conjunction with other factors such as helicases and ssDNA-binding proteins that enhance its processivity. Additionally, a FEN activity is required in HR pathways, particularly in single-strand annealing (SSA) and the resolution of D-loops. Therefore, OEX1 is a primary candidate for resolving such recombination intermediates.

The FEN activity of OEX1 is likely also essential for BER, which is a principal mechanism for repairing damaged mtDNA in both plants and animals (Szczesny et al., 2008; Boesch et al., 2011; Ferrando et al., 2018). Okazaki fragment processing and long-patch BER share similar mechanisms, as both involve a DNA polymerase displacing the existing strand to form a flap, which is then cleaved by a FEN. Similar to BER, the FEN activity of OEX1 might also play a role in ribonucleotide excision repair (RER), a process that removes ribonucleotides misincorporated into DNA during DNA replication. RER requires an RNase H to cleave the strand containing the ribonucleotide, and the strand-displacement activity of a DNA polymerase to create a flap, which is then processed by a FEN, such as OEX1.

Finally, OEX1 might also play a role in the removal of R-loops, which can form in transcribed regions and are known to cause genomic instability (Gan et al., 2011; Wimberly et al., 2013). This conclusion is supported both by the strong *in vitro* activity of OEX1 in degrading the RNA strand of RNA:DNA hybrids, and by the DRIP-qPCR results that revealed accumulation of R-loops in the mitogenome of *oex1-1,* particularly in the highly transcribed regions encoding the rRNAs.

### OEX1 is critical for plant development and its absence causes mtDNA instability

The phenotype of *oex1-1* plants confirmed the importance of OEX1 in plant growth, development, and fertility. By the third generation of homozygous mutants, plants could no longer produce offspring. This quasi-sterility can be attributed to severely deformed pistils and stigmas, along with reduced pollen viability, as evidenced by failed attempts to pollinate wild-type pistils with *oex1-1* pollen. Although the overall phenotypic defects were qualitatively consistent among individual plants, their severity varied and correlated with the degree of mitochondrial genome instability. Typically, this instability resulted in changes in the relative abundances of mitochondrial genomic regions, as arising from preferential amplification of sub-genomes created by ectopic recombination on small repeats (in the range of 100-500 bp). However, as seen in some *oex1-1* plants, much smaller and imperfect repeats as well as microhomologies could also drive ectopic recombination in the mutant. These sub-genomic mtDNA molecules may result from the accumulation of unprocessed DNA ends in the *oex1-1* mutant, thus triggering strand invasion and replication re-initiation via break-induced replication (BIR) or microhomology-mediated BIR (MMBIR). As previously suggested, processes that generate mtDNA breaks potentially promote error-prone repair pathways, and likely explain the mtDNA instability and increased heteroplasmy observed in recombination surveillance mutants (Marechal and Brisson, 2010; Gualberto and Newton, 2017).

Whether the severity of the *oex1-1* phenotypes is directly linked to the stoichiometric changes of a specific mitochondrial locus, is currently unknown. However, although changes in relative copy numbers of mtDNA regions can influence the mitochondrial transcriptome (Wallet et al., 2015), the regions with reduced copy numbers in *oex1-1* plants do not contain any known genes. Furthermore, our transcriptomic data revealed no potentially detrimental changes in the expression or processing of mitochondrial gene transcripts. On the contrary, several gene transcripts were more abundant, possibly due to an increased copy number of their corresponding coding sequences.

Thus, the harmful effects of the *oex1* mutation on plant development is likely due to problems in mtDNA replication and/or the aberrant segregation of recombination-generated subgenomes. These defects seem to affect mitochondrial biogenesis, though not necessarily through obvious problems with gene expression. As revealed by expression of a promoter:GUS fusion, *OEX1* is primarily expressed in young, rapidly dividing tissues, consistent with the observed defects in mitochondrial biogenesis and proliferation revealed by TEM imaging. Similar effects on mitochondrial size have also been observed in mutants deficient in the mitochondrial ssDNA-binding protein OSB1 (Zaegel et al., 2006). This suggests a possible link between mtDNA replication and/or segregation and mitochondrial division, and is reminiscent of the situation in mammalian cells, where nucleoids involved in active mtDNA replication localize to contact points with the endoplasmic reticulum, which form the sites of mitochondrial division (Lewis et al., 2016). However, such a functional connection remains to be demonstrated in plants. An additional component potentially contributing to the developmental defects in mutants with impaired mtDNA maintenance are retrograde responses that activate (or inactivate) the expression of nuclear genes and, in this way, suppress cell proliferation. Such responses have been observed in animal cells, and in plants, in chloroplast signaling and in the *radA* mutant (Tigano et al., 2021; Chevigny et al., 2022; Fu et al., 2023).

### Differential roles for the two OEX1 isoforms generated by alternative splicing

Both our experimental data and publicly available transcriptomic datasets confirm that Arabidopsis has two OEX1 isoforms that are encoded by alternatively spliced transcripts. These isoforms seem conserved among flowering plants, as corresponding variants have been found in other species including monocots, thus pointing to their biological relevance. The extension found in mitochondrial OEX1a is neither found in chloroplast OEX2 nor in the bacterial orthologs, thus raising the question why mtDNA maintenance requires two isoforms. Our expression analysis suggests developmental regulation, with a higher OEX1b/OEX1a ratio in developing flowers. We also demonstrated that both proteins are functional, as they complement the *oex1-1* developmental defects. However, they influence mtDNA segregation differently, and plants complemented with OEX1b maintained a slight imbalance in mtDNA subgenomes, which, by contrast, was fully corrected in plants complemented with OEX1a. This difference may suggest distinct roles or substrate preferences of the two isoforms. Structural modeling supports this assumption, by placing the OEX1a extension near the active site and the DNA-binding domain. Indeed, *in vitro* activity tests showed that, while both isoforms process the same substrates, they differ in processivity, with OEX1a having a higher specific activity on all tested substrates, including 5’-3’ exonuclease activity on DNA ends and flap endonuclease (FEN) activity. Nevertheless, we cannot rule out the alternative possibility that OEX1b is specifically required to process certain specific recombination intermediates. The existence of two isoforms might reflect distinct interactions with other factors involved in mtDNA maintenance. OEX1 localization in nucleoids suggests that the protein may be part of larger nucleo-protein structures where mtDNA metabolism occurs. The requirement of DNA polymerase activity for efficient processing of DNA gaps, such as long-patch BER intermediates, supports the idea that OEX1a and OEX1b act in concert with other factors, possibly by forming transient interactions. Interestingly, Y2H analyses revealed that the two isoforms interact differently with the organellar DNA polymerases POL1A and POL1B. While both isoforms are able to interact with POL1A, only OEX1b interacts with POL1B. POL1A and POL1B have distinct roles in mtDNA maintenance. While they are redundant for genome replication, they differ in processivity, fidelity and involvement in DNA repair (Parent et al., 2011; Baruch-Torres and Brieba, 2017; Ayala-Garcia et al., 2018; Garcia-Medel et al., 2019; Schatz-Daas et al., 2022). Our findings from plants complemented with either OEX1a or OEX1b under genotoxic stress provide additional support for the idea that the two OEX1 isoforms have partially differing roles in mtDNA repair.

The picture emerging from this work and from other studies is that mtDNA maintenance relies on a fine-tuned interplay of multiple factors. These include RecA recombinases, SSB and OSB ssDNA-binding proteins, DNA polymerases, and OEX nucleases, which are present in different isoforms with specialized roles shaped by tissue-specific expression, activity profiles, and distinct functions in replication and repair pathways. These isoforms can arise from distinct paralogous genes, dual targeting of the same protein to both organelles, or, in the case of OEX1, alternative splicing.

## Materials and Methods

### Plant material and constructions

Sequence data from this article can be found in the Arabidopsis Genome Initiative or GenBank/EMBL databases under the following accession numbers: OEX1, At3g52050; OEX2, At1g34380. *Arabidopsis thaliana* T-DNA insertion line *oex1-1* (GK-911E05) derives from accession Col-0 and was obtained from the Nottingham Arabidopsis Stock Centre (NASC). Plants were grown on soil or *in vitro* under a 16 h light/8 h dark photoperiod (65-85 μmol photons m^−2^ s^−1^) at 21 °C/18 °C. For *in vitro* cultures, surface-sterilized seeds were sown on agar plates containing half-strength MS255 media (Duchefa) supplemented with 1% (w/v) sucrose and stratified 2 days at 4 °C in the dark. For DNA and RNA extraction, plant tissue frozen in liquid nitrogen was ground with a TissueLyser II (Quiagen). DNA was extracted using the cetyltrimethylammonium bromide method and RNA was extracted using TRI Reagent (Molecular Research Centre, Inc.). RNA was further extracted with phenol-chloroform before ethanol precipitation. For mutant complementation, the WT *OEX1* gene and promoter sequences were cloned into binary vector pGWB613 and used to transform heterozygous *oex1-1* plants. For complementation with either OEX1a or OEX1b sequences, the gene sequence including the promoter, 5’ UTR, and the first two exons and introns was fused to the *OEX1a* or *OEX1b* cDNA sequences. For the hemicomplementation constructs, the organellar targeting sequence (OTS) of OEX1 (86 codons) was replaced with either the OTS (49 codons) of mitochondrial alternative oxidase 1 from soybean (AOX1; NM_001249237) or the OTS (80 codons) of the chloroplast small subunit of ribulose bisphosphate carboxylase (RBCS; At1g67090). The constructs were assembled using MultiSite Gateway cloning in vector pB7m34GW (https://gatewayvectors.vib.be/), giving AOX1:OEX1 and RBCS:OEX1 constructs. These were used for *Agrobacterium tumefaciens* transformation of heterozygous *oex1-1* plants by floral dip.

### Phylogenetic Analysis

For the building of phylogenetic trees, bacterial and plant orthologous sequences were aligned with T-Coffe implemented in the MacVector package with default parameters (Myers-Miller dynamic algorithm, gap open penalty of -50 and gap extension penalty of -50) and a consensus maximum likelihood tree was built with IQ-TREE (http://iqtree.cibiv.univie.ac.at), from 1000 bootstrap iterations. Graphical representation and edition of the tree was with TreeDyn (v198.3).

The Arabidopsis OEX1 sequence was modelled on the structure of the 5’-3’ exonuclease domain of *Thermus aquaticus* DNA Polymerase I (Kim et al., 1995), using Modeler (Sali and Blundell, 1993); http://salilab.org/modeller/about_modeller.html), and the modeled structures represented with PyMol (https://pymol.org/2/).

### Subcellular localization and Promoter-GUS fusion

For *in vivo* intracellular localization, the *OEX1* and *OEX2* gene sequences were cloned into the pUBC-paGFP-Dest vector, upstream and in frame with the GFP coding sequence, under the control of the UBQ10 promoter (Grefen et al., 2010). Leaf pavement cells of independent transformants were observed at the confocal microscope (Zeiss LSM700). For colocalization of OEX1 with mitochondria, the *OEX1* cDNA was amplified and cloned into the pUBC-mCherry-Dest binary vector (Grefen et al., 2010), to obtain the pUbi::OEX1-mCherry construct. The resulting plasmid was used for Agrobacterium-mediated transformation of plants expressing the mitochondrial marker pCaMV35S::mTP-GFP, corresponding to the transit peptide of ATP citrate lyase (At2g20420) fused to GFP, under the control of the 35S promoter. The resulting seeds were selected on MS medium supplemented with phosphinotricin at a final concentration of 10 mg/L. This observation was facilitated by examining the leaves of plants grown under conditions that induce elongated mitochondria, as described (Jaipargas et al., 2015). To facilitate visualization of mitochondrial nucleoids, cells were observed under conditions that induce elongated mitochondria, as described (Jaipargas et al., 2015). To this end, Arabidopsis seedlings expressing both pCaMV35S::mTP-GFP and pUbi::OEX1-mCherry transgenes were grown in the dark for 8 days on 0.5x MS medium without sugar. DNA was stained with DAPI solution (1 µg/mL) in phosphate-buffered saline for 30 min at 25 °C in the dark, and leaf pavement cells were imaged at a Leica TCS SP8 confocal laser-scanning microscope. Excitation of DAPI was at 405 nm, GFP and chlorophyll at 488 nm and mCherry at 561 nm. Fluorescence emission was collected at 435– 475 nm for DAPI, 500–545 nm for GFP, 595-645 nm for mCherry and beyond 650 nm for chlorophyll.

For promoter-GUS histochemical analysis, the 5’ upstream region of OEX1 (595 nucleotides comprising 161 nucleotides of *OEX1* 5’-UTR and a large part of upstream gene At3g52040) was cloned upstream of the *GUS* gene in the binary vector pMDC162 (Curtis and Grossniklaus, 2003), and the construct was used to generate stable Arabidopsis transformants. Tissues from six independent lines were stained with 5-bromo-4-chloro-3-indolyl-ß-D-glucuronic acid (X-Gluc; Biosynth) and observed by stereomicroscopy and transmission microscopy.

### Transmission Electron Microscopy

Leaf tissue samples were fixed overnight in 3% glutaraldehyde, treated 2 h with 10% (w/v) picric acid, 2 h with 2% uranyl acetate and stained with 0.1% (v/v) osmium tetroxide in 150 mM phosphate buffer, pH 7.2. After dehydration through an ethanol series, samples were infiltrated with EPON812 medium-grade resin (Polysciences). Ultrathin 70 µm sections were collected on grids coated with formvar (Electron Microscopy Sciences) and visualized with a Hitachi H-600 electron microscope operating at 75 kV.

### Yeast two-hybrid assays

The cDNA sequences of Arabidopsis OEX1a, OEX1b, POL1A and POL1B were cloned into pGADgwT7-AD and pGBKgwT7-BD vectors. The yeast strain AH109 was co-transformed with different combinations of AD and BD vectors and transformants were selected on synthetic defined (SD)/-Leu/-Trp (SD-LW) medium for 4 days at 30 °C. Colonies were grown overnight in SD-LW liquid media, and after adjustment to OD_600nm_= 0.5, serial dilutions were plated on SD-LW and on SD/-Leu/-Trp/-His (SD-LWH) plates. The interaction between the F-box protein EBF2 and transcription factor EIN3 was used as a positive control.

### qPCR / RT-qPCR Analysis

For RT-qPCR experiments, 5 μg of RNA were depleted from contaminating DNA by treatment with RQ1 RNase-free DNase (Promega) and reverse-transcribed with Superscript IV Reverse Transcriptase (Thermo Fisher Scientific), according to the manufacturer’s protocol using random hexamers. The qPCR assays were performed with the LightCycler480 (Roche) on a reaction mix of 6 µL containing 1X LightCycler 480 SYBRGreen I MasterMix (Roche) and 0.5 μM of each primer. Three technical replicates were performed for each experiment and Cp values were determined from the second derivative maximum. Quantification of mtDNA and cpDNA copy numbers used a set of primer pairs located along the organellar genomes, as described previously (Wallet et al., 2015; Chevigny et al., 2022). Results were normalized against *UBQ10* (At4g05320) and *ACT1* (At2g37620) nuclear genes. The accumulation of ectopic recombination in mtDNA was quantified using primers flanking each repeat, as described (Miller-Messmer et al., 2012), and results normalized against the *COX2* (AtMG00160) and 18S rRNA (AtMG01390) mitochondrial genes. RT-qPCR experiments were normalized using the *GAPDH* (At1g13440) and *ACT2* (At3g18780) transcripts as standards.

### Sequencing

Total leaf DNA of WT and *oex1*-*1* plants was quantified with a QuBit Fluorometer (Life Technologies) and libraries prepared with the Nextera Flex Library kit according to the manufacturer instructions (Illumina), using 100 ng DNA. Final libraries were quantified, quality checked on a Bioanalyzer 2100 (Agilent) and sequenced on an Illumina Miseq system (2 × 150 paired-end reads). For RNAseq, total RNA was extracted from WT and *oex1-1* 10 day-old seedlings, tree biological replicates each, using NucleoSpin® RNA Plant kit (Macherey-Nagel). RNAs were quantified by QuBit (Invitrogen) and quality checked on Bioanalyzer 2100. Five micrograms of total RNAs were ribodepleted using riboPOOL kit (siTOOLs Biotech). Ten nanograms of ribodepleted RNAs were used for library preparation with NEBNext Ultra II Directional RNA Library Prep, according to the manufacturer instructions. Libraries were sequenced (1 × 50 bases single read) on an Illumina NextSeq^TM^ 2000. DNAseq and RNAseq reads were mapped to the mtDNA with Bowtie2, implemented in the MacVector Assembler package application. Because of sequences of chloroplast origin inserted in the mtDNA, reads mapping to the chloroplast genome were first removed from the analysis. Coverage profiles were generated with the MacVector Assembler package.

### Expression and purification of recombinant proteins

The coding sequence of the At*OEX1* gene minus the first 85 codons was amplified from cDNA and both alternative cDNA sequences, *OEX1a and OEX1b,* were cloned in the pET28a expression vector (Novagen), to express recombinant proteins containing a N-terminal hexa-histidine tag. The recombinant proteins were soluble and were purified from the *E. coli* BL21(DE3) extracts by nickel ion affinity chromatography, followed by gel filtration (Figure S7). Briefly, protein expression was induced with 0.5 mM IPTG and after 2 h at 37 °C the bacterial cells were sedimented, resuspended in 150 mM Tris-HCl pH 8.0, 5% glycerol, 150 mM NaCl (Buffer A) supplemented with 1 mM PMSF, and lysed with a LM20 Digital Microfluidizer® Processor (Microfluidics™) under 1200 PSI. The crude lysate was clarified by centrifugation at 17 000 g for 20 min, adjusted to 30 mM imidazole. Proteins were purified by affinity on cOmplete™ His-Tag Purification Resin (Roche®). Columns were washed with Buffer A followed by washing with 50 mM imidazole. The recombinant proteins were eluted with a 90-310 mM imidazole step gradient, followed by gel filtration on a Superdex 200 10/300 GL column equilibrated with Buffer A (Figure S5C). Protein aliquots were flash frozen in liquid nitrogen and stored at -80 °C. A catalytic mutant protein was equally expressed and purified, which contained a substitution of an essential aspartic acid with alanine in the active site (protein D250A, Figure S5B). Following purification, the recombinant proteins were free of detectable contaminants and purified as a single monodispersed peak by gel filtration, of a size consistent with a monomeric protein (Figure S7C). The optimal activity conditions of OEX1 on 5’-labeled dsDNA were tested (Figure S8). The protein requires Mg^2+^ ions, that could be poorly substituted by Mn^2+^, but not by Zn^2+^ or Ca^2+^. It showed optimal activity at about 37 °C but was inhibited at higher temperature. Finally, it worked in a broad pH range, between pH 7.4 and pH 9.0. Based on these results we established the assay conditions for further experiments as 2 mM Mg^2+^, pH 8.0 at 37 °C.

### Nuclease assays

Structure-specific nuclease assays were performed using oligonucleotides synthesized by Integrated DNA Technologies (IDT). DNA oligonucleotides were either radiolabelled in 5’ using T4 polynucleotide kinase (Fermentas) and [γ-^32^P]ATP, or in 3’, by template-directed addition of one to three cytosines by the Klenow fragment of *Escherichia coli* DNA polymerase I and [α-^32^P]dCTP. RNA oligonucleotides were labeled in 3’ using poly(U) polymerase (NEB) and [α-^32^P]UTP. When necessary, labeled products where annealed with cold oligonucleotides by incubation in 10 mM Tris-HCl pH 7.5, 1 mM EDTA, 100 mM NaCl (Buffer B) during 5 min at 95 °C, followed by slow cooling to room temperature. Oligonucleotide sequences used to form the different structures are listed in supplementary Table 1. Substrates were purified by electrophoresis on non-denaturing 15% polyacrylamide gels and eluted in Buffer B, overnight at 4 °C under agitation. Standard nuclease reactions were performed with 50 fmol of labeled structure in reaction buffer (50 mM Tris-HCl pH 8.0, 1 mM DTT, 0.1 mg/mL BSA and 4% glycerol) supplemented with 2 mM bivalent ion and 1.5 µM enzyme. The reaction was incubated at 37 °C and stopped by addition of an equal volume of stop-mix (95% formamide, 20 mM EDTA, 0.05% bromophenol blue, 0.05% xylene cyanol). Reaction products were analysed by denaturing PAGE (17% 19:1 acrylamide/bisacrylamide, 7 M urea, 1 X TBE) and revealed using an Amersham Typhoon biomolecular imager (GE Healthcare Life Sciences).

### DNA:RNA Immunoprecipitation

Total DNA was extracted with Plant II Midi kit (Macherey-Nagel®) and fragmented with EcoRI, BamHI, HindIII and XbaI to cut the mtDNA into pieces between 200 to 5,000 bp. About 5 µg were either treated or not with Ribonuclease H (RNase H 5 U/μL) (Thermo Fisher Scientific®), and incubated with 2 µL S9.6 antibody (Merck) specific for RNA:DNA hybrids, for 15 h at 4 °C under rotation (10 rpm), followed by incubation with 25 µL of Protein G Sepharose beads (Sigma-Aldrich®) for 2 h at 4 °C (Yang et al., 2017). Immunoprecipitated fragments were detached from antibodies by proteinase K treatment (Thermo Fisher Scientific®) and purified by phenol/chloroform extraction followed by ethanol precipitation. The isolated DNA was analysed by qPCR using primer pairs targeting the highly transcribed mtDNA regions coding for the ribosomal RNAs, and results were normalized against the quantification of non-transcribed regions of the mtDNA.

## Supplemental Figures

**Supplemental Figure S1. Widespread expression of *oex1*, according to promoter-GUS fusion.**

Promoter::GUS fusion reporter plants of OEX1 (pOEX1::GUS) were stained for GUS gene expression at different developmental stages. A) three-day-old seedling, B) root, C) 10-day-old seedling, D) inflorescences, E) flower, and F) developing silique.

**Supplemental Figure S2. Knockout of *OEX1* in the oex1-1 line.**

RNAseq sequence coverage of the *OEX1* gene in WT and *oex1-1,* showing the last six exons are no longer transcribed in the T-DNA insertion mutant line. The vertical arrow indicates the position of the T-DNA insertion, at the border between the 9^th^ exon and 9^th^ intron.

**Supplemental Figure S3. Transmission electron microscope images showing smaller, more electron-dense mitochondria in *oex1-1* compared to WT.**

**A)** Cells from leaves of same size showed morphologically normal chloroplasts (cp) in *oex1,* while mitochondria (mt) were smaller and more electron-dense as compared to mitochondria in WT cells. **B)** Distribution of the organelle sections (in µm^2^) extracted from TEM images. Asterisks indicate statistical significance (p<0.001) according to Student’s t-test.

**Supplemental Figure S4. Alternative splicing results in the expression of two OEX1 isoforms.**

**A)** Alternatively spliced exon 6 of OEX1 encodes an 18 aa insertion that is absent from the sequences of OEX2 and the 5’-3’ exonuclease of bacterial DNA Pol I. An insertion of comparable size is found in the OEX1 sequences of land plants. **B)** Comparison between the known structure of the 5’-3’ exonuclease domain of DNA Pol I from *Thermophilus aquaticus* and the modeled structure of OEX1. The 18 aa extension is in green and indicated by an arrow, with the H3TH domain in purple. The two essential aspartic acids in the PIN active domain are highlighted in red. **C)** Relative quantification of alternative spliced *OEX1a* and *OEX1b* transcripts in different tissues.

**Supplemental Figure S5. Interaction between OEX1 and organellar DNA polymerases.**

**A)** Interaction between OEX1a and OEX1b with the organellar DNA polymerase POL1A, tested by yeast two-hybrid assay. BD and AD are the binding and activating domains of GAL4, respectively. Serial dilutions of cell suspensions were spotted on SD-LWH plates. The interaction between EBF2 and EIN3 served as positive control. Both BD-OEX1a and BD-OEX1b could interact with AD-POL1A. **B)** As in A, but with organellar DNA polymerases POL1B. BD-OEX1b could interact with AD-POL1B, but not BD-OEX1a. Controls with empty vectors confirmed no significant autoactivation by either BD-OEX1a/1b or by AD-POL1A/1B. **C)** Data from the complexome database (complexomemap.de/at_mito_leaves) indicate that OEX1 migrates as a band smaller than 100 kDa on Blue-Native gels. For comparison, the results obtained for the TWINKLE helicase, subunit B2 of gyrase and ssDNA-binding proteins SSB1 and SSB2 are also shown.

**Supplemental Figure S6. RNAseq analysis of the mitochondrial transcriptome**

Comparison of the coverage profiles of mtDNA transcripts reveals no significant differences in the processing of protein-coding gene transcripts between *oex1-1* and WT. Variations in the coverage profiles for rRNAs are attributed to differences in ribodepletion efficiency.

**Supplemental Figure S7. Expression and purification of recombinant OEX1 proteins.**

**A)** SDS-PAGE analysis of the expression and solubility of OEX1a and OEX1b, indicated by arrowheads. **B)** expression of a catalytic mutant where an essential aspartic acid of the catalytic PIN domain was replaced by an alanine. **C)** Purification of recombinant protein OEX1a by gel filtration, following metal-affinity purification on a Ni-column. The apparent molecular weight is consistent with a monomeric protein.

**Supplemental Figure S8. Optimization of OEX1 activity conditions.**

**A)** Activity of OEX1a on a 5’-labeled dsDNA substrate, tested at varying concentrations of divalent ions. Substrates and degradation products were analyzed on denaturing polyacrylamide gels, after 3 minutes of incubation. Maximum activity was observed with Mg^2+^. **B)** As in A, according to incubation temperature. The optimal temperature observed was 37 °C. **C)** As in A, but according to pH. The optimum observed was pH 8.0.

**Supplemental Figure S9. Sequence-dependent processivity and activity on ssDNA and DNA gaps.**

**A)** Activity of OEX1a on dsDNA substrates differing by 1, 2 or 3 consecutive Gs, showing pausing at G positions. **B)** Little activity of OEX1a and OEX1b on ssDNA substrates. **C)** Minimal activity of OEX1 alone on DNA gap of 1, 3 or 10 nucleotides, but no detectable activity of the non-catalytic mutant (D250A).ss

