## Supplementary figures and images for "R-Loop control and mitochondria genome stability require the 5’-3’ exonuclease/flap endonuclease OEX1"

### Supplemental Figures

**Figure S1.**

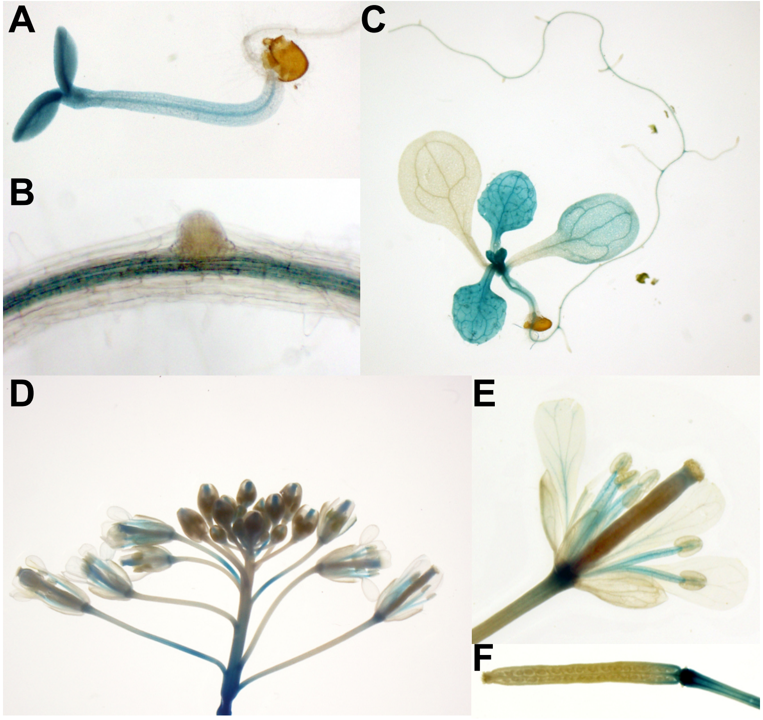

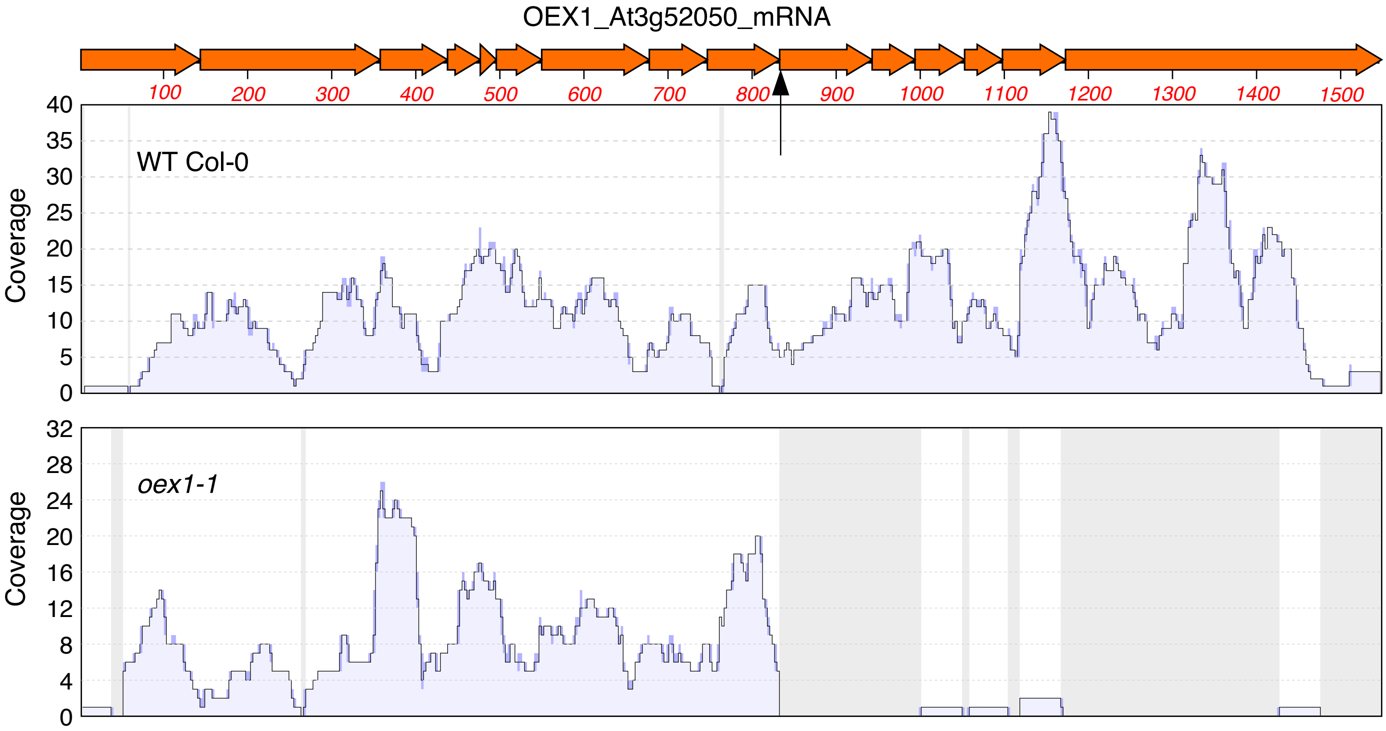

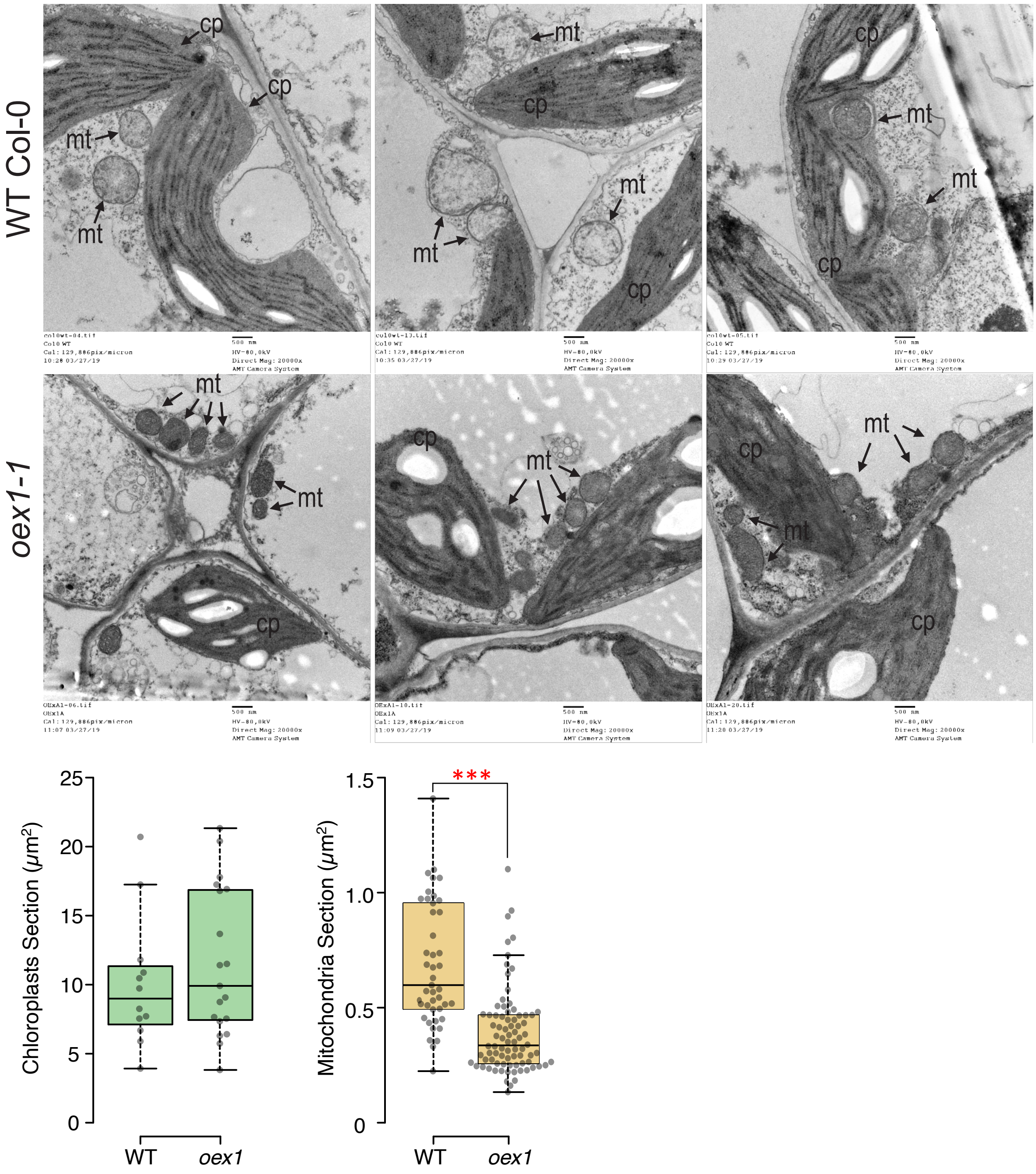

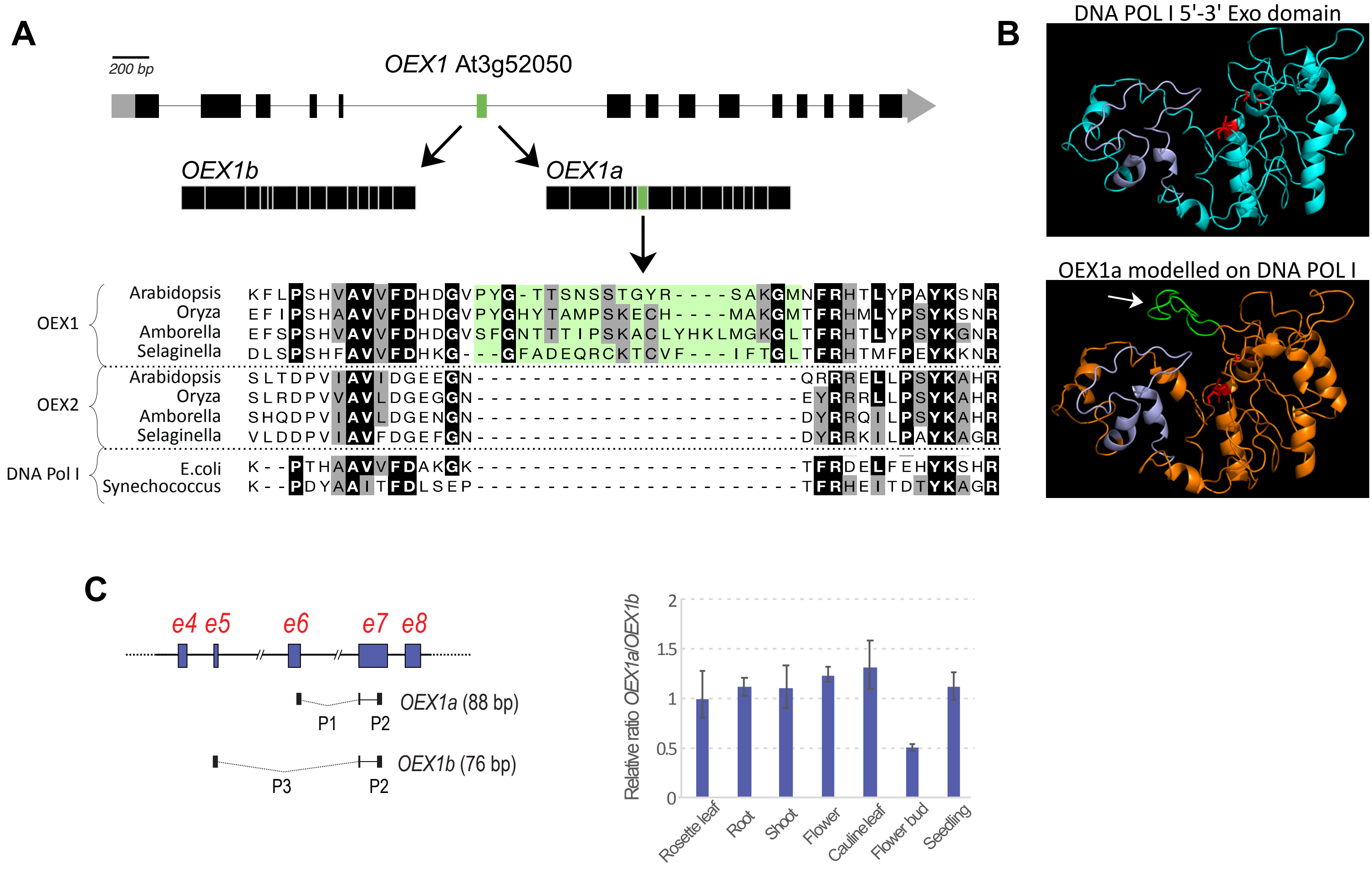

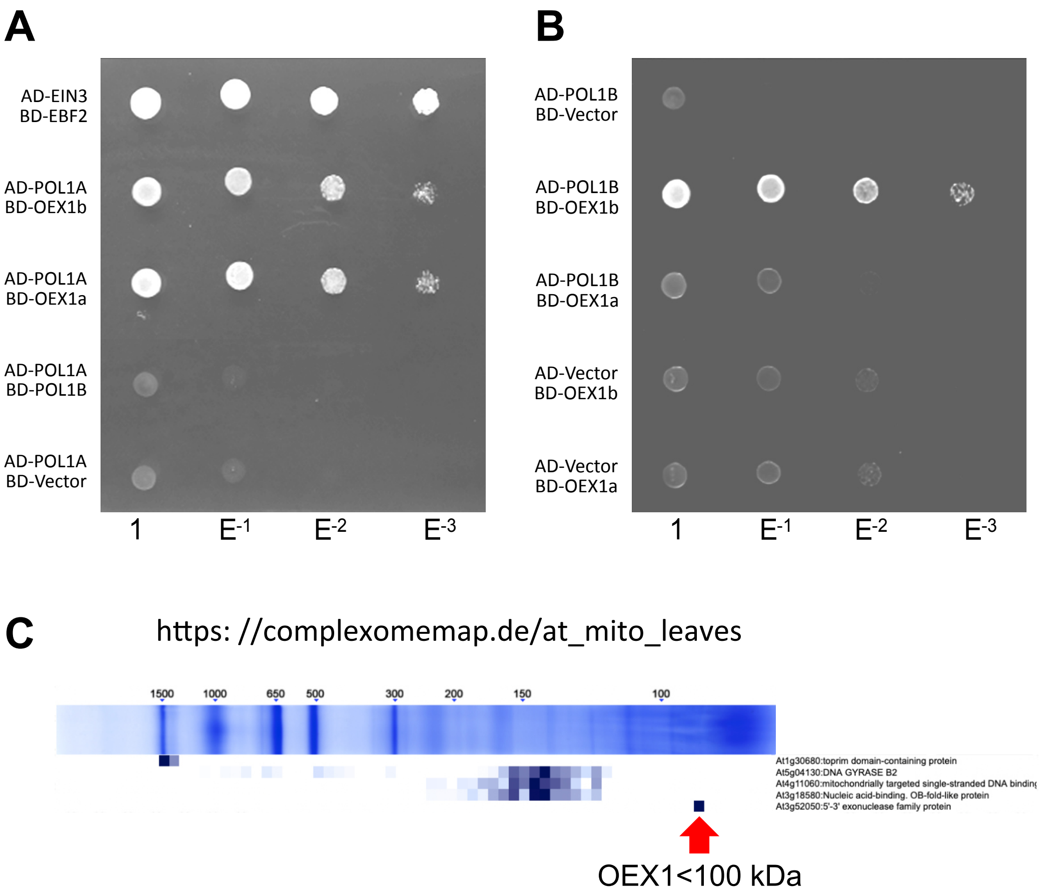

Figure S6.

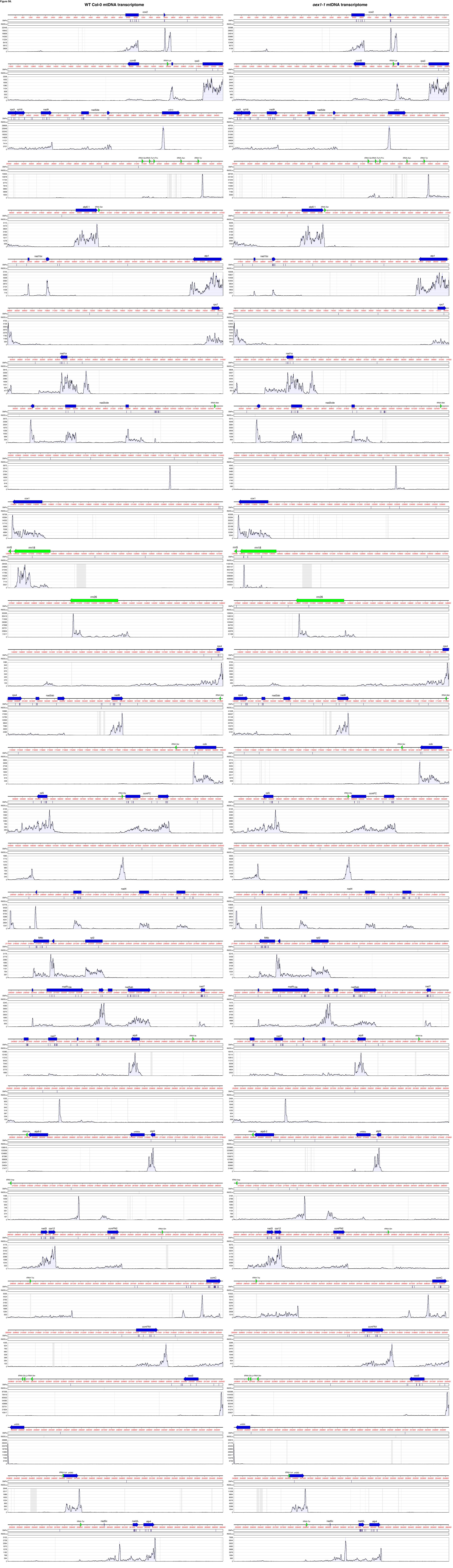

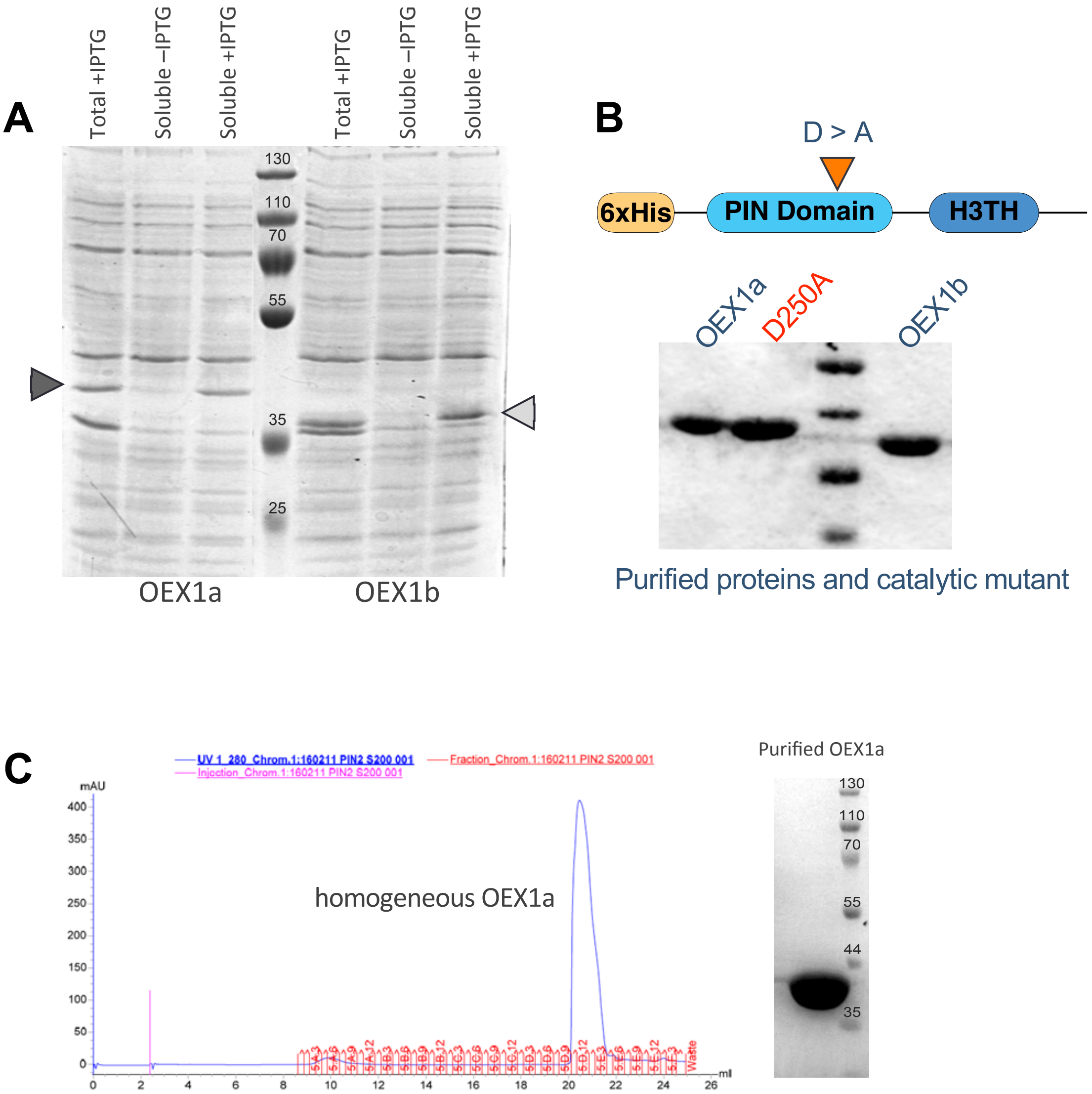

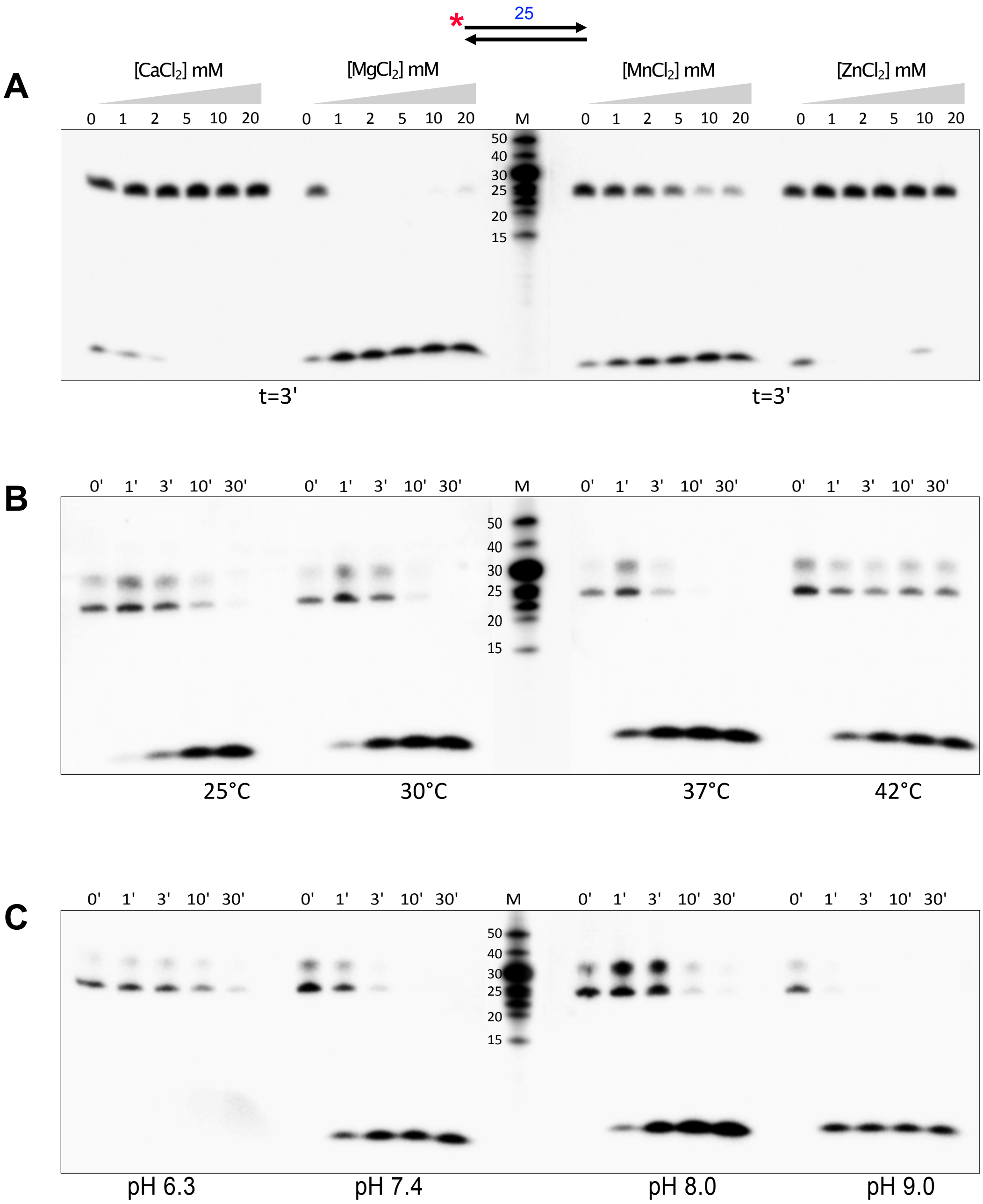

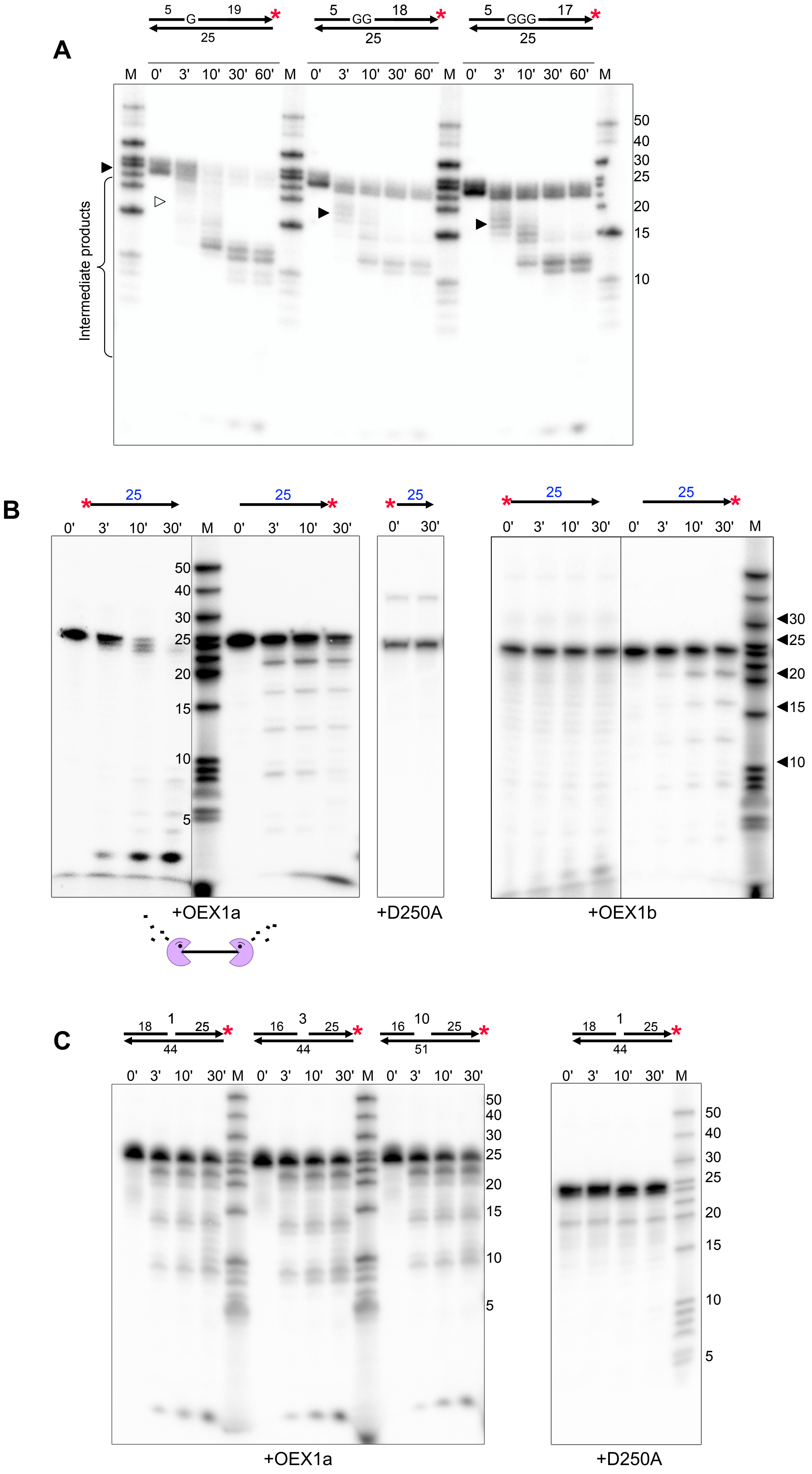
